# A unified atlas of CD8 T cell dysfunctional states in cancer and infection

**DOI:** 10.1101/2020.07.25.220673

**Authors:** Yuri Pritykin, Joris van der Veeken, Allison Pine, Yi Zhong, Merve Sahin, Linas Mazutis, Dana Pe’er, Alexander Rudensky, Christina Leslie

## Abstract

CD8 T cells play an essential role in defense against viral and bacterial infections and in tumor immunity. Deciphering T cell loss of functionality is complicated by the conspicuous heterogeneity of CD8 T cell states described across different experimental and clinical settings. By carrying out a unified analysis of over 300 ATAC-seq and RNA-seq experiments from twelve independent studies of CD8 T cell dysfunction in cancer and infection we defined a shared differentiation trajectory towards terminal dysfunction and its underlying transcriptional drivers and revealed a universal early bifurcation of functional and dysfunctional T cell activation states across models. Experimental dissection of acute and chronic viral infection using scATAC-seq and allele-specific scRNA-seq identified state-specific transcription factors and captured the early emergence of highly similar TCF1^+^ progenitor-like populations at an early branch point, at which epigenetic features of functional and dysfunctional T cells diverge. Our atlas of CD8 T cell states will facilitate mechanistic studies of T cell immunity and translational efforts.

## INTRODUCTION

Upon completing their differentiation in the thymus, mature naïve T lymphocytes enter the periphery and recirculate through secondary lymphoid organs, where, upon an encounter with a cognate antigen in the presence of co-stimulatory molecules, they become activated, expand, and differentiate into effector or memory T cells. These cells then take up residence in lymphoid and non-lymphoid organs where they exert their immune functions. In contrast, chronic or suboptimal antigenic stimulation, e.g. in the absence of co-stimulation, can result in a state of hypo-responsiveness or anergy (1). Over the last decade, this simple textbook view has evolved into a markedly more nuanced and complex picture of T cell differentiation with a plethora of seemingly distinct states emerging from a large number of studies in mice and man (2–4). CD8 T cells, whose function is essential for defense against viral and bacterial infections and for tumor immunity, serve as a case study in this regard. Phenotypic and functional analyses of CD8 T cells in acute and chronic viral infections, cancer, transplantation and “self” tolerance in both experimental animal models and in human patients have offered numerous descriptions of activated effector, long-lived central and short-lived effector memory cells and their precursors, as well as an array of CD8 T cell states with perturbed functionalities dubbed anergic, exhausted, and reversibly or irreversibly dysfunctional (5–20). Recent characterization of a small subset of exhausted/dysfunctional cells, named “stem cell-like” or progenitor cells, capable of self-renewal, adds further complexity to the topography of CD8 T cell activation and differentiation (6, 7, 18–26). T cell dysfunction therefore presents a challenging case study for resolving gene regulatory programs and differentiation trajectories towards distinct cellular fates.

Studies of CD8 T cell dysfunctional states, besides being highly significant for understanding of basic mechanisms of adaptive immunity, have attracted particular attention due to the realization that prevention or reversal of CD8 T cell dysfunction can serve as a potent strategy for the treatment of both solid organ and hematologic malignancies and chronic infections. Inefficient mobilization of endogenous CD8 T cell responses or a failure to engage them in cancer patients in response to checkpoint blockade inhibitors, as well as disease resistance or relapse following adoptive CD8 T cell therapies including CAR (chimeric antigen receptor)-T cells, have been attributed to a large degree to dysfunction or exhaustion of tumor- and virus-specific CD8 T cells (26–28). Transcriptional and chromatin features associated with these states have been extensively explored through the analyses of epigenomes and transcriptomes of isolated subsets of functional and dysfunctional CD8 T cells using DHS-, ATAC-, and RNA-seq and through single cell transcriptomics and proteomics (9–20, 25, 29–31). These studies have significantly advanced the knowledge of CD8 T cell differentiation and highlighted pronounced changes in T cell chromatin states. However, the remarkable heterogeneity of functional and dysfunctional states of CD8 T cells revealed by these genome-wide analyses in diverse experimental and clinical settings poses a major problem of distinguishing between common vs. context-specific features of CD8 T cell differentiation towards dysfunction and the underlying regulatory mechanisms. For example, in chronic viral infection, T cells encounter antigen in an inflammatory and stimulatory context and have been described as progressing through an effector state prior to differentiating to an exhausted state (32–34). Meanwhile, a two-step process of differentiation from a reversible to an irreversible dysfunctional state was reported in the setting of early tumorigenesis, where naïve tumor-specific T cells encounter antigen in a non-inflammatory setting that may result in inadequate priming or activation (12, 32). Therefore, it is currently unclear how to reconcile these different models of progression to dysfunction, and the differentiation programs giving rise to two distinct states at late time points of chronic antigen exposure – self-renewing dysfunctional progenitors and terminally exhausted cells – are not well understood.

A vexing obstacle in addressing these issues has been the inability to directly compare genome-wide data from different studies due to technical sources of variation, including but not limited to sample preparation, sequencing quality, batch effects, and cell numbers, making meaningful integration of massive amounts of data generated in mouse and human studies problematic. To address this challenge, we carried out a uniform reprocessing and a statistically principled batch effect correction approach to over 300 chromatin accessibility (ATAC-seq) and gene expression (RNA-seq) datasets generated in twelve independent studies of CD8 T cell states observed across experimental mouse models of acute and chronic infection and tumor models. Our analysis revealed a universal signature of chromatin accessibility changes in the progression to terminal dysfunction in both tumors and chronic infection, implying early commitment to a dysfunctional fate in all settings of chronic antigen exposure. The chromatin state observed at early time points during the development of dysfunction was similar to that of dysfunctional progenitor cells found in late time points in infection and tumor models. Motif-based regression modeling of this unified chromatin accessibility compendium enabled inference of state-specific transcription factor activities and implicated new factors in the progression to terminal dysfunction. This bulk-level T cell state analysis suggested a universal early bifurcation of functional and dysfunctional T cell activation states across models of cancer and chronic infection.

We further characterized this early branch point by carrying out single-cell analysis of CD8 T cell populations in the context of acute and chronic lymphocytic choriomeningitis virus (LCMV) infection and observed a TCF1^+^ progenitor-like population resembling the memory precursor effector cell (MPEC) population in acute infection (8, 35, 36). Regression modeling of scATAC-seq clusters allowed us to refine the association of T cell functional states with activity of transcription factors, whose causal role was established through a comprehensive scRNA-seq atlas in T cell populations from F1 hybrid mice combining evolutionary distant genomes of laboratory and wild-derived mouse strains.

Together, these results provide new insights into the early emergence of a progenitor-like population in response to chronic antigen stimulation that appears to precede the establishment of dysfunction-committed progenitor cells. Our unified atlas of CD8 T cell chromatin and expression states across mouse models and single-cell analyses of the bifurcation between functional and dysfunctional T cell responses will provide a valuable resource to the community and facilitate further mechanistic and translational studies.

## RESULTS

### Dysfunctional T cells in tumors and chronic viral infection share a common epigenetic and transcriptional state space

We collected 166 chromatin accessibility (ATAC-seq) datasets (**Figure 1A**) from 10 recent studies on CD8 T cell function in mouse models of infection and cancer. These included T cells of various antigen specificities profiled in acute bacterial infection, acute and chronic viral infection, including memory precursor cells and tissue-resident memory cells, as well as tumor-infiltrating lymphocytes in hepatocarcinoma and melanoma models. The data included adoptively transferred endogenous and CAR-T cells; T cells with and without treatment with anti-PD1 immunotherapy; and progenitor/stem cell-like T cell populations isolated from multiple models of chronic antigen stimulation (9–16, 18, 35). We performed a uniform processing of these data sets to construct a high-resolution atlas of 129,799 reproducible chromatin accessibility peaks across CD8 T cell states; these peaks were further split into 221,054 subpeaks and associated to genes (**Methods**).

**Figure 1.**
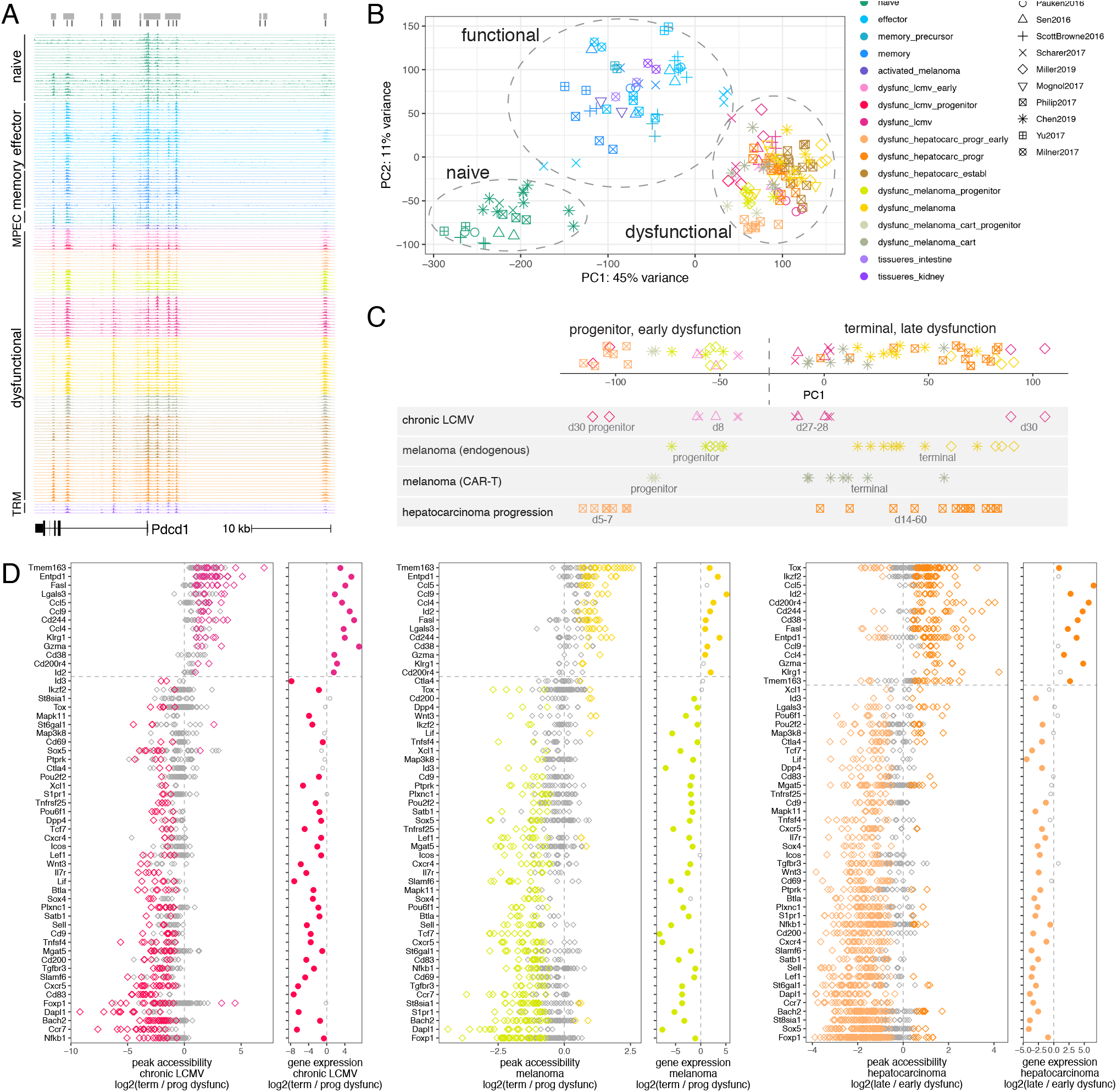
Dysfunctional CD8 T cells in tumors and chronic viral infection share a common chromatin state space. **A.** Snapshot of the ATAC-seq compendium near the *Pdcd1* locus. **B.** Principal component analysis (PCA) of library-size normalized batch-effect corrected ATAC-seq read counts in 150bp windows around peak summits (functional cell state shown by color, data source by symbol). **C.** First principal component (PC1) of PCA for library-size normalized batch-effect corrected ATAC-seq read counts in differentially accessible peaks in dysfunctional T cells from different studies. For clarity, vertical random jiggle is added, and subsets of samples from different studies are shown below. **D.** In each panel, left: library-size normalized batch-effect corrected ATAC-seq read count log2 fold change for peaks (diamond shown in color for significantly decreased/increased, FDR < 0.05) of significantly differentially accessible genes; right: log2 fold change of RNA-seq gene expression for the same genes. Shown are comparisons between progenitor and terminally dysfunctional T cells in chronic LCMV infection and in melanoma, and between early and late states of dysfunction in hepatocarcinoma progression.

Expectedly, PCA analysis of ATAC-seq read counts in the peaks showed that the samples clustered by data source (**Supplementary Figure 1A**). We therefore applied a generalized linear model (GLM) accounting for the sample’s data source and functional state (naïve, functional, dysfunctional, see **Methods**). After this batch effect correction, distances between functionally related samples from different studies (**Supplementary Figure 1B**) decreased, and samples readily clustered into broad functional categories regardless of the data source (**Figure 1B**). A more conservative GLM correction not explicitly accounting for the differences between functional and dysfunctional cells produced similar results (**Supplementary Figure 1C**). Furthermore, naïve cells from multiple studies clustered together in the PCA plot, while effector and memory cells (as well as pre-activated cells injected in melanoma-bearing mice, but ignorant of the tumor antigen) formed a cluster of functional cells. Interestingly, cells profiled at day 4 in acute infection were positioned between clusters of naïve and functional cells, suggesting their intermediate state. Strikingly, the chromatin states of dysfunctional tumor-infiltrating T cells from different tumor models and T cells in chronic viral infection formed a distinct cluster, suggesting there exists a universal program of T cell dysfunction observed across models and immune challenges. Focusing on the most significantly differentially accessible peaks between functional and dysfunctional cells again confirmed the similarity between T cell chromatin states in tumors and chronic infection, distinct from naïve and functional T cells (**Supplementary Figure 1D,E**). A similar analysis of 136 samples of gene expression (RNA-seq) data from eight studies (9, 11–14, 17–19) showed consistent results (**Supplementary Figure 1F**). Differential expression was significantly correlated with differential accessibility (**Supplementary Figure 1G**). This confirmed that dysfunctional T cells are epigenetically and transcriptionally similar in the settings of chronic infection and across tumor models.

Genes with the strongest differential accessibility between functional and dysfunctional cells at their promoter, intronic, and nearby intergenic peaks (**Methods**) included well-known markers of T cell activation, cytotoxicity, adhesion, and apoptosis, as well as those encoding key transcription factors, cytokines and cytokine receptors, and other cell surface and intracellular proteins (**Supplementary Figure 2**). Consistent with the overall correlation between differential accessibility and differential expression (**Supplementary Figure 1G**), in many cases (but not always) differential accessibility of individual genes was associated with significant differential expression (**Supplementary Figure 2**). This integrative differential accessibility and differential expression analysis across our compendium allowed us to identify a high confidence universal epigenetic and transcriptional gene signature of T cell dysfunction.

### T cell temporal progression in both tumors and chronic viral infection mirrors the state change from progenitor to terminal dysfunction

Dysfunctional T cells profiled as early as at day 5-8 after antigen encounter in tumor or chronic viral infection models clustered closely with the terminally dysfunctional cells profiled at day 22-35 rather than with effector or memory cells (**Figure 1B**, **Supplementary Figure 1F**). Cells characterized as progenitor/stem-cell-like dysfunctional cells in different studies also clustered with the terminally dysfunctional cells. This suggested that T cells differentiate towards dysfunction early after antigen encounter rather than first mounting and then losing a functional effector response, in contrast to proposed exhaustion models in chronic viral infection (32–34). To further understand this differentiation process and compare it across models, we next focused on chromatin accessibility changes in dysfunctional cells.

We first compared genome-wide accessibility profiles of different subsets of dysfunctional T cells across models and immune challenges. We observed that peaks differentially accessible between progenitor and terminally dysfunctional (“exhausted”) cells, including endogenous or transferred CAR-T cells in melanoma or dysfunctional cells in chronic viral infection, were consistent between models and studies and also displayed chromatin accessibility changes between early (d5-8) and late (d22-35) states of dysfunction in chronic viral infection and hepatocarcinoma tumorigenesis (**Figure 1C**, **Supplementary Figure 3A-C**). Furthermore, changes in accessibility between early and late dysfunctional states, as well as between progenitor and terminally dysfunctional states, correlated with changes in gene expression, suggesting a role of chromatin accessibility in regulating gene expression changes (**Supplementary Figure 3D**). Importantly, T cells at early time points in all mouse models were similar in bulk chromatin state to dysfunctional progenitor cells, a subpopulation sorted from late (“exhausted”) time points in chronic viral infection or tumors. Differential accessibility between progenitor and terminally dysfunctional CD8 T cells in the melanoma mouse model was significantly associated with that in TILs from melanoma patients (36), suggesting generalizability of the analysis in mice to human patients (**Supplementary Fig 3E**). This analysis of genome-wide chromatin accessibility profiles suggested that there exists a common axis of differentiation of T cell dysfunction observed across models and immune challenges.

We also found that many individual genes were consistent across models in their pattern of expression and chromatin accessibility along this common differentiation axis. Many of the same genes were differentially accessible in progenitor and terminally dysfunctional cells in chronic viral infection (**Supplementary Figure 4A**) and melanoma (**Supplementary Figure 4B**), and in early and late dysfunction states in chronic viral infection (**Supplementary Figure 4C**) and hepatocarcinoma (**Supplementary Figure 4D**), including numerous transcription factors and genes associated with T cell function (**Figure 1D**). For example, genes encoding markers of terminal dysfunction Entpd1 (Cd39), 2B4 (Cd244), Cd38 were significantly more accessible in terminal exhaustion and late tumor-specific dysfunction, while genes encoding markers of progenitor cells Cxcr5, Slamf6, IL7Ra chain were significantly more accessible in progenitor and early dysfunctional cells. Additional cell surface proteins encoded by genes significantly more accessible and expressed in progenitor cells, such as CD9, CD200, TNFRSF25 (DR3), CD83, CD69, could also serve as candidate markers for progenitor cells. Loci encoding chemokines Ccl4, Ccl5, Ccl9 were more accessible in the terminal dysfunctional state, consistent with its characterization as hyperactivated. The locus of the transcription factor Id2 was more accessible in terminal/late dysfunction, while Id3, Tcf7, Lef1, Nfkb1, Pou2f2, Pou6f1 were more accessible in progenitor or early dysfunctional cells, suggesting the involvement of multiple transcription factors in the establishment and maintenance of these states. In most cases, differential accessibility at individual genes in early/progenitor versus late/exhausted states was associated with differential expression, which was consistent with overall correlation between differential expression and differential accessibility (**Supplementary Figure 3D**). *Tox* and *Ikzf2*, whose heightened expression was previously demonstrated to be associated with terminal dysfunction, were indeed significantly more accessible and highly expressed in terminal vs. early T cell dysfunction during hepatocarcinoma tumorigenesis; however, both loci had many peaks significantly more accessible in progenitor cells and Ikzf2 was significantly overexpressed in progenitor cells both in chronic LCMV infection and melanoma, suggesting the active role of these transcription factors in the progenitor cells (**Figure 1D**). Thus, our analysis identified universal markers and transcription factors associated with T cell differentiation towards dysfunction across multiple models and organisms.

### Coordinated activity of many transcription factors is associated with functional and dysfunctional T cell states

We next sought to identify what transcription factors (TFs) are associated with different T cell activation and differentiation states, particularly the progenitor cells. We analyzed TF targets, as predicted by motifs of sequence-specific TF binding, and performed a supervised modeling of chromatin accessibility data based on TF motif sites to infer state-specific TF activities. For each sample, we used negative binomial regression with ridge regularization to predict absolute levels of chromatin accessibility (ATAC-seq read coverage) in peaks from TF motif occurrences (**Figure 2A**, **Supplementary Figure 5A**, **Methods**). For this analysis, we focused on 105 motifs for TFs expressed in CD8 T cells after grouping TFs with indistinguishable motifs. Using coefficients of the regression models, we estimated the effect on accessibility of each TF in each sample. This allowed us to map chromatin accessibility profiles into a lower dimensional inferred transcription factor activity space, largely preserving the relationships between samples (**Supplementary Figure 5B**).

**Figure 2.**
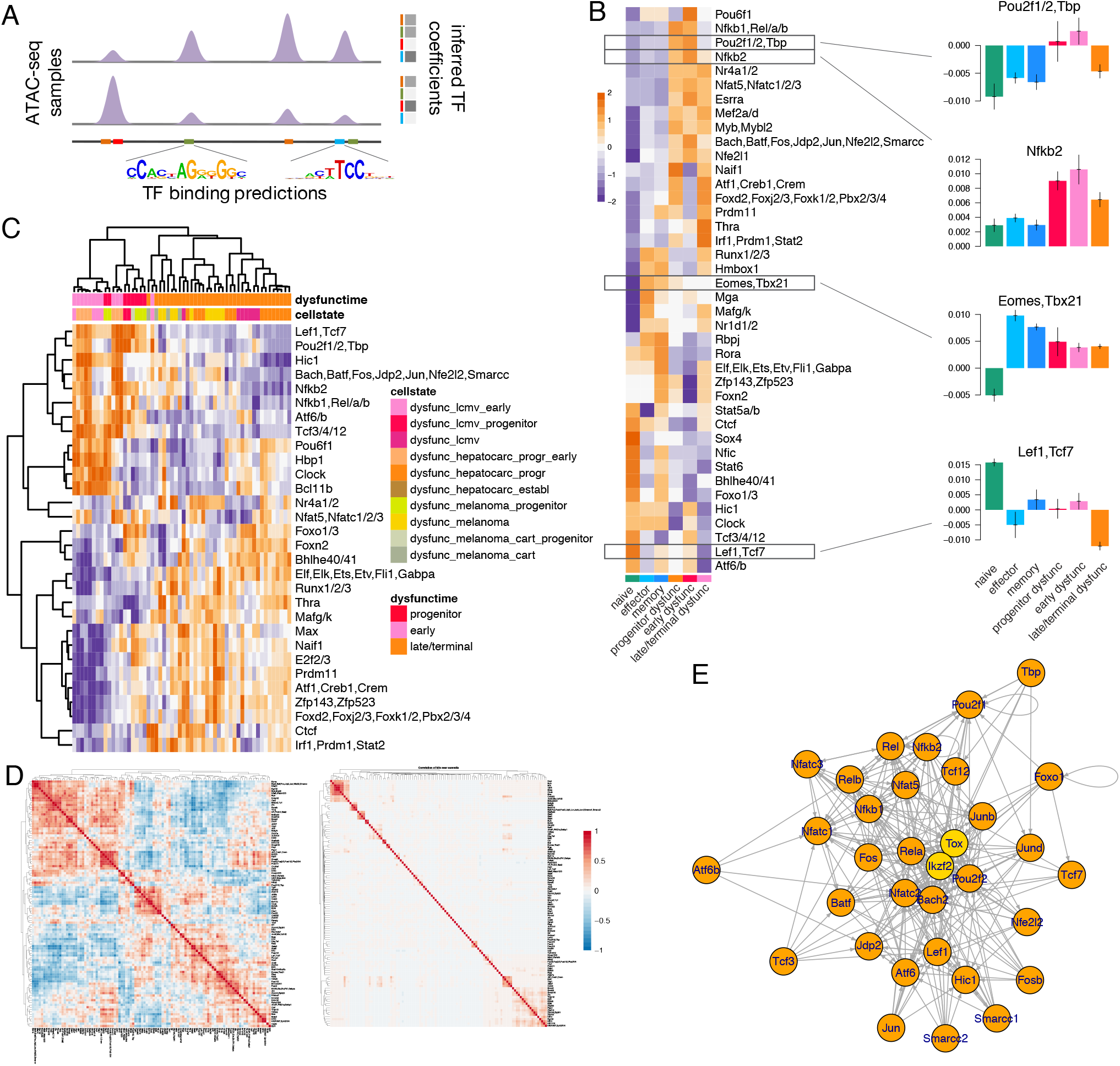
Predictive modeling identifies transcription factors associated with progenitor dysfunctional CD8 T cells. **A.** Schematic of the negative binomial generalized linear regression analysis to infer transcription factor (TF) associations with chromatin accessibility. **B.** Inferred TF motif coefficients from the regression model of library-size normalized batch-effect corrected ATAC-seq signal in each sample, consolidated between replicates of the indicated conditions across studies. Inferred coefficients with the highest variance across conditions are shown (z-score row normalized). **C.** Inferred TF motif coefficients with the highest variance across dysfunctional states (z-score row normalized). **D.** Spearman correlation of inferred TF motif coefficients across all antigen-experienced samples (left) and of TF motif match scores across the atlas of peak summit regions (right). **E.** Network of predicted regulatory interactions between TFs in progenitor dysfunctional T cells. Binding motifs are not available for TOX and IKZF2 (nodes shown in yellow).

Next, in order to identify TFs most strongly associated with T cell functional states, we consolidated the most variable inferred transcription factor activities across replicates and studies (**Figure 2B**). For example, as expected, the Eomes/Tbx21 motif was associated with low accessibility in naïve cells and high accessibility in all antigen-experienced cells, while the Lef1/Tcf7 motif was associated with high accessibility in naïve cells and low accessibility in terminally dysfunctional cells (**Figure 2B**). Motifs of Nr4a, Nfkb, Nfat, Pou, AP1 families were strongly associated with high accessibility in early dysfunctional and progenitor cells as opposed to terminally dysfunctional, naïve or functional cells, while motifs of Zfp143 and Zfp523 were associated with high accessibility in terminal dysfunction (**Figure 2B,C**); this included both previously (11–14, 19, 21, 33) and newly identified factors. Interestingly, Ctcf motif binding was associated with differential accessibility between conditions (**Figure 2B,C**), suggesting potential differences in chromatin looping between T cell functional states (37, 38); consistently, we observed a significant relationship between differential accessibility and differential expression when associating ATAC-seq peaks to genes using chromatin loops defined using published Hi-C data in naïve CD8 T cells (39) instead of our default approach (**Supplementary Figure 5C**, **Methods**). Overall, this motif-based regression modeling of ATAC-seq data was a powerful approach for prioritizing candidate TFs that could be associated with T cell functional states and conditions.

To gain global understanding of TF regulatory programs across T cell states, we next looked at the overall correlation of inferred TF activities across all antigen-experienced cells. We found multiple clusters of correlated TF activities, potentially suggesting coordinated regulatory programs in different functional states (**Figure 2D**). Importantly, correlated TFs in the same cluster for the most part had different motifs and were bound at different sites (**Figure 2D**). Focusing on the progenitor dysfunctional cells, we found that many TFs expressed in this condition had multiple predicted binding sites in loci encoding other TFs and thus could potentially regulate their expression (**Figure 2E**). This suggested that the coordinated and hierarchically organized activity of a broad range of TFs, not necessarily binding at the same sites, may be required for establishing and maintaining T cell functional states, as opposed to a single “master regulator” or handful of TFs.

### Single-cell chromatin accessibility analysis reveals the early emergence of a progenitor-like T cell population

Our integrative analysis of bulk ATAC-seq data across mouse models found an early (d7-8) divergence of CD8 T cells between responses to acute and chronic immune challenges (**Figure 1B**) and identified potential driver TFs at cell population level. Furthermore, plastic and reprogrammable cells early (d7-8) in the development of T cell dysfunction were more similar to immunotherapy-responsive progenitor dysfunctional cells identified at later (d20-35) time points than to terminally dysfunctional cells (**Figure 1C**). Therefore we decided to better characterize chromatin states of the early divergence between functional and dysfunctional fates and explain the similarity to the progenitor population at the single-cell level. Furthermore, we wanted to obtain a higher resolution map of TF regulators of T cell functional states and confirm our predictions from analysis of sorted cell populations. To that end, we performed single-cell chromatin accessibility (scATAC-seq) analysis of the total splenic CD8 T cell compartment in mice at day 7 (d7) upon infection with LCMV Armstrong (Arm), resulting in acute infection, and LCMV clone 13 (Cl13), resulting in chronic infection (**Figure 3A**).

**Figure 3.**
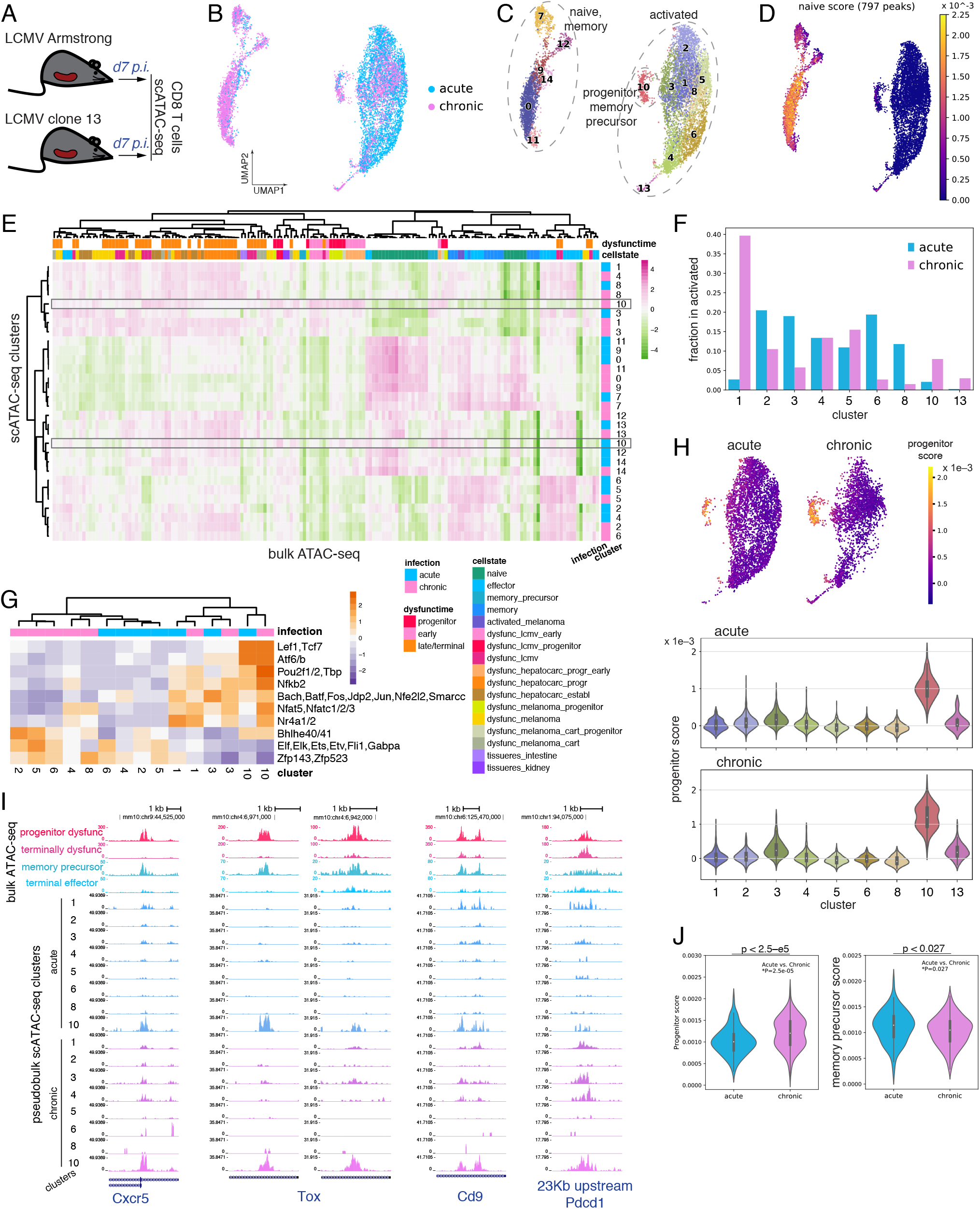
scATAC-seq analysis reveals heterogeneity and overlapping but divergent CD8 T cell responses to acute and chronic immune challenges. **A.** Experimental setup. **B.** UMAP representation of TF-IDF-transformed scATAC-seq data. **C.** Clustering of scATAC-seq data using the Louvain algorithm. **D.** Single-cell heatmap showing the naïve cell signature derived from bulk ATAC-seq data. **E.** Association by ssGSEA of library-size normalized batch-effect corrected bulk ATAC-seq data with library-size normalized scATAC-seq counts aggregated over cells in each cluster in each of the two samples (z-score row normalized). **F.** Barplot showing the fraction of cells in each sample from clusters 1-8, 10, 13 that belong to each cluster. **G.** Inferred TF motif coefficients for scATAC-seq counts averaged over cells in each cluster (for clusters 1-8, 10, 13) in each of the two samples. Inferred coefficients with the highest variance are shown (z-score row normalized). **H.** Progenitor dysfunctional signature derived from bulk ATAC-seq data scored in scATAC-seq data (for cells in clusters 1-8, 10, 13 only). Top: single-cell heatmap of the signature score, separately for acute and chronic infection. Bottom: violin plot of the signature score for cells in each cluster, separately in acute and chronic infection. **I.** Genome browser tracks of ATAC-seq data for selected peaks. Bulk ATAC-seq for progenitor and terminally dysfunctional cells, and for terminal effector and memory precursor cells. Normalized aggregated single-cell ATAC-seq for cells in each cluster (for clusters 1-8, 10) in each of the two samples. **J.** Progenitor dysfunctional cell and memory precursor cell signatures derived from bulk ATAC-seq data scored in cells from cluster 10 in acute and chronic infection (Mann-Whitney U test).

We first constructed a combined atlas of 189,281 chromatin accessibility peaks that included both peaks identified from the bulk ATAC-seq data compendium and 59,482 newly identified scATAC-seq peaks. Quantification and filtering yielded 4,767 and 5,865 cells, with 12,598 and 13,195 median read count per cell, initially isolated from Arm and Cl13 infected mice, respectively. Normalization using term frequency–inverse document frequency (TFIDF), dimensionality reduction using PCA followed by UMAP, and Louvain clustering (**Methods**) suggested that d7 responses to infection with the two LCMV clones were heterogeneous and overlapping with each other (**Figure 3B,C**, **Supplementary Figure 6A,B**).

Next we sought to characterize functional features of cells captured with scATAC-seq. For this, we used ssGSEA (40) to associate scATAC-seq cell clusters with the bulk ATAC-seq data compendium, and epigenetic signatures derived from bulk ATAC-seq data (**Figure 3D,E**, **Supplementary Figure 6C,D**, **Methods**). We found that cells in clusters 0, 7, 9, 11, 12, 14 were likely a mixture of naïve and background-activated memory cells that were not specific to and therefore did not respond to the LCMV antigen; on the other hand, clusters 1-6, 8, 10, 13 likely consisted of LCMV responding cells. While both responses were heterogeneous and highly overlapping, we also found differences in responses. For example, cluster 1 was most similar to dysfunctional cells and was dominated by cells from Cl13 infected mice, while cluster 6 was most similar to effector cells and dominated by cells from Arm (**Figure 3E,F**, **Supplementary Figure 6E**). This suggested a clear divergence between the CD8 T cell responses to Arm and Cl13 viruses as early as d7 post infection.

To validate and refine our predictions of cell-state specific TF activities for bulk cell populations (**Figure 2, Figure 1D**), we applied the same motif-based predictive modeling to pseudo-bulk chromatin accessibility signal aggregated from cells within each scATAC-seq cluster, separately for acute and chronic infection. This allowed us to identify TFs most strongly associated with different functional states in scATAC-seq data (**Figure 3G**, **Supplementary Figure 6F**). These results were largely consistent with regression analysis in bulk data but helped to identify more precisely which TFs are associated with different functional states of T cells early in response to acute or chronic infection.

Interestingly, cells in cluster 10, when compared with the bulk ATAC-seq data, were most similar to the progenitor dysfunctional cells profiled at late chronic infection (**Figure 3E,H, Methods**). This suggested that not only in established chronic infection by Cl13, but also at d7 of infection, cells in a progenitor-like dysfunctional state were observed. Furthermore, surprisingly, cluster 10 also contained a small number of cells from acute infection that were similar to memory precursor cells (8, 11, 16, 41, 42) based on comparison with the bulk ATAC-seq data (**Figure 3F,H**, **Supplementary Figures 6E, 7A,B**). These cluster 10 cells from acute and chronic infection were similar in their genome-wide chromatin accessibility profiles (and thus belonged to the same cluster in scATAC-seq data), in their association with TF activities, including enrichment of Lef1/Tcf7, Nr4a, Nfkb, Nfat, Pou, AP1 family motifs (**Figure 3G**), and also in the chromatin accessibility levels at gene loci encoding proteins important for T cell activation and function, including progenitor marker Cxcr5 (**Figure 3I**, **Supplementary Figure 7A,B**), consistent with our observations from bulk ATAC-seq data analysis in progenitor dysfunctional cells (**Figure 1E**, **Figure 2B,C**). This shared cluster suggested that progenitor-like cells, by analogy to memory precursor cells, are established as early as at d7, and may give rise to more differentiated cell states as previously reported for progenitor dysfunctional cells from later time points (18, 20, 21). However, we also observed evidence of differential accessibility between the progenitor-like and precursor cells in chronic and acute infection (**Figure 3J**, **Supplementary Figure 7C**) and potentially different TF activity (**Figure 3G**), suggesting an even earlier divergence between functional and dysfunctional fates.

By complementing our bulk ATAC-seq analysis across models, the scATAC-seq analysis provided an epigenetic-level characterization of the early divergence between CD8 T cell responses to acute and chronic immune challenges, identified a progenitor-like subpopulation common to but also likely divergent between acute and chronic infection, and implicated cell-state specific TFs.

### Single-cell transcriptional analysis confirms a progenitor/precursor T cell population in early response to both acute and chronic infection

To further explore the heterogeneity and temporal progression of CD8 T cell responses at single-cell resolution, to better understand the relationship between T cell chromatin states and phenotypes, and to validate our state-specific TF predictions through allele-specific analyses, we performed single-cell RNA-seq analysis of CD8 T cells from hybrid mice generated upon breeding C57Bl/6J and SPRET/EiJ mice [F1(B6xSpret)] at three time points during acute and chronic LCMV infection (**Figure 4A**, **Supplementary Figure 8**). This approach takes advantage of combining in F1 T cells wide-spread natural genetic variation (~40 million SNPs) between the evolutionary distant parental genomes and assessing its effect on gene expression and chromatin accessibility. Virus-specific cells were profiled before infection (d0) and upon activation at day 7 and day 40 (d40) during the acute LCMV infection and activated CD62L-negative cells were profiled at day 7 and day 35 (d35) during the chronic LCMV infection. Cells isolated from chronically infected B6 mice at d35 were used as a control. Bulk RNA-seq and ATAC-seq profiling of CD8 T cells at d0, d7, and d60 upon acute LCMV infection in F1(B6xSpret) mice (43) confirmed their consistency with the counterparts from B6 mice (**Supplementary Figure 9A**).

**Figure 4.**
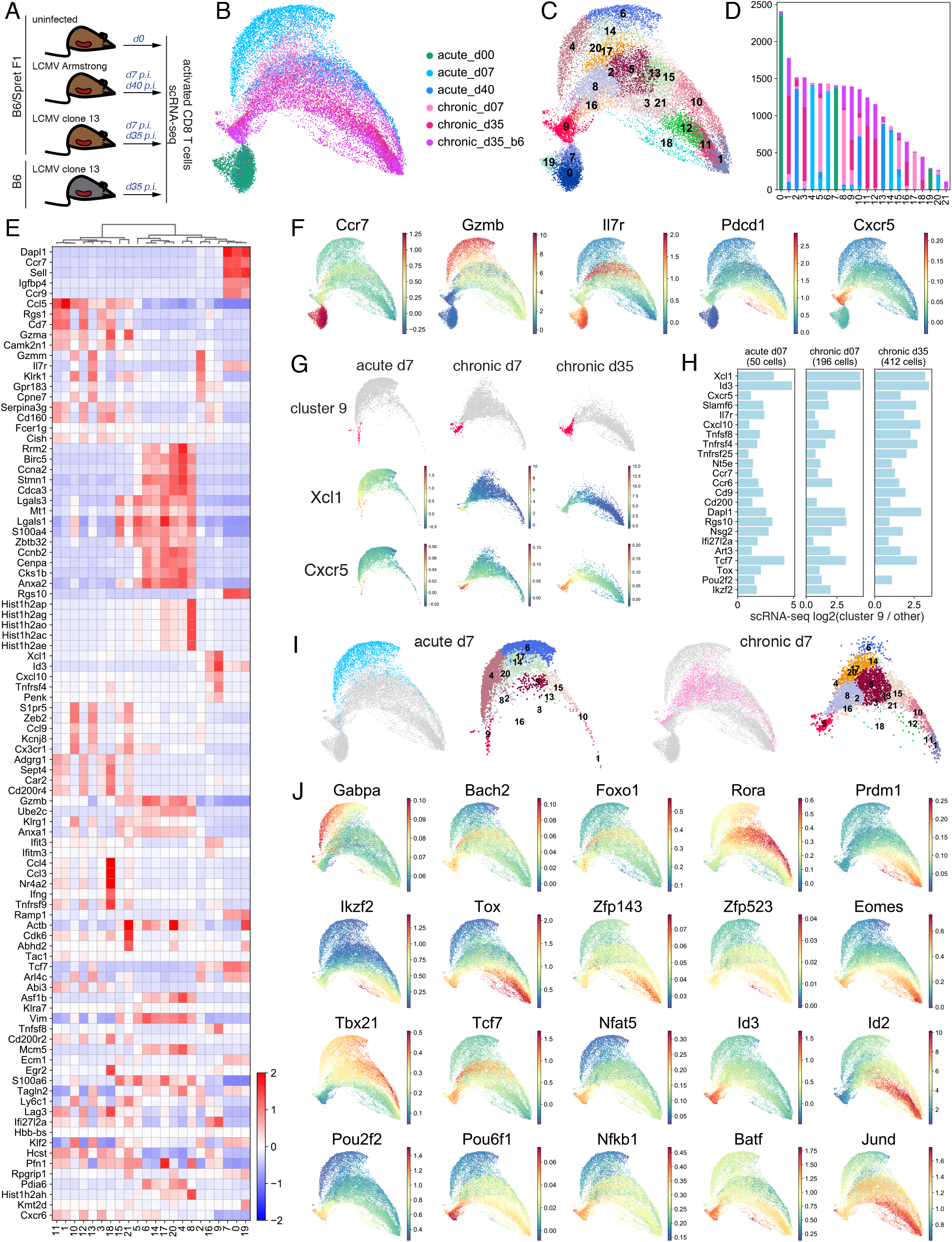
scRNA-seq analysis uncovers phenotypic heterogeneity of CD8 T cell response and cell-state specific transcription factor expression. **A.** Experimental setup. **B.** UMAP representation of library-size normalized scRNA-seq data. **C.** Clustering of scRNA-seq data using the Louvain algorithm. **D.** Barplot showing the number of cells in each cluster from each sample. **E.** Average scRNA-seq gene expression (library-size normalized UMI counts) in each cluster for differentially expressed genes between clusters (z-score row normalized). **F.** MAGIC-imputed gene expression for selected genes. **G.** MAGIC-imputed gene expression in individual samples. **H.** Log2 fold change of expression of selected genes in cells from cluster 9 vs. all other cells in individual samples. **I.** Cells at d7 in acute and chronic infection separately and their cluster composition, within overall UMAP. **J.** MAGIC-imputed expression of selected genes encoding transcription factors.

After scRNA-seq filtering steps (**Methods**), our data set consisted of 9,822 genes profiled in 24,400 cells, ranging from 3,419 to 4,824 cells per sample, with median 1,147 expressed genes per cell. UMAP embedding revealed a well-separated naïve state and diverging phenotypic arms in acute and chronic responses with heterogeneity within each sample and gradients of expression of well-known T cell function markers potentially reflecting differentiation trajectories (**Figure 4B**, **Supplementary Figure 9B,C**). Louvain clustering identified clusters that for the most part had representation from multiple samples, suggesting the different time points and immune challenges included cell populations in similar functional states, albeit in different proportions (**Figure 4C,D**). Cells at d35 in chronic infection from B6 and F1(B6xSpret) mice aligned on top of each other, appeared in similar proportions in most clusters and were closest in the kNN graph, suggesting a similar overall T cell response in the two models (**Figure 4B-D**, **Supplementary Figure 9D,E**). Dimensionality reduction using other well-known methods (tSNE, force-directed layout, diffusion map) produced similar results (**Supplementary Figure 9F-H**).

We used a combination of complementary approaches in order to functionalize the scRNA-seq data. First, unbiased differential expression analysis identified genes enriched in each cluster (**Figure 4E**, **Supplementary Figure 10A,B**, **Methods**). Second, visualizing the expression of the T cell signature genes (**Figure 4F**, **Supplementary Figure 9C, 10C**) further helped to interpret the clusters and characterize subsets of cells. Clusters 0, 7, 19, dominated by cells profiled before infection (**Supplementary Figure 10D**), were enriched for naïve T cell marker genes Ccr7, Ccr9, Lef1, Tcf7. Gzmb was highly expressed in clusters 4, 6, 14, all dominated by cells at d7 upon acute infection, consistent with effector function. Cell proliferation and survival related genes Birc5, Stmn1, Cdca3 were enriched in clusters 4, 6, 8, 14, 17, 20, which were dominated by cells from early response (d7) to acute or chronic infection. Clusters 2, 10, 13 displayed high Il7r expression and were dominated by cells from acute infection at d40, which were mostly memory cells. Clusters 1, 3, 11, 12, 13, 21 were dominated by cells at d35 in chronic infection and enriched for markers of dysfunction Pdcd1, Lag3, Entpd1, and Cd38. Clusters 5, 15 were dominated by cells from chronic infection at d7 and enriched for Lgals1, Lgals3, Mt1, Mt2. Cluster 18 consisted of dysfunctional cells enriched for Pdcd1, Lag3, and Entpd1 that also overexpressed Ifng, Ccl3, Ccl4, and Nr4a2 and thus were also highly activated. Clusters 9 and 16 overexpressed markers of progenitor cells Cxcr5, Slamf6, Tcf7, and Id3. Differential expression analysis restricted to cells in each sample relative to the same clusters confirmed that their functional characterization was correct (**Supplementary Figure 11**). Finally, association of our bulk RNA-seq compendium (**Supplementary Figure 1F**) with scRNA-seq cluster-specific expression profiles using ssGSEA largely confirmed our cluster characterizations (**Supplementary Figure 12, Methods**).

Thus, our analysis identified cell clusters encompassing archetypal CD8 T cell subsets including naïve T cells, memory precursor cells, short-lived effector cells, central memory, effector memory, terminally dysfunctional cells and their precursors. This cellular spectrum is similar to but more detailed than previously published scRNA-seq datasets of CD8 T cell responses in acute or chronic infection (18, 30, 31, 44, 45), suggesting that these cellular states are consistently generated across independent studies, and using mice of distinct genetic backgrounds.

We expected that our scRNA-seq profiling would be able to capture progenitor dysfunctional and progenitor-like cells. Indeed, consistent with our scATAC-seq analysis (**Figure 3**), our scRNA-seq analysis identified a subpopulation of cells (cluster 9) (**Figure 4G**) resembling the previously characterized progenitor/memory-like/stem-cell-like dysfunctional cells based on the enrichment of signature genes such as Id3, Tcf7, Slamf6, and Cxcr5 (**Figure 4E,F**, **Supplementary Figures 10A, 13A**) and the comparison with bulk RNA-seq from sorted subpopulations (**Supplementary Figure 12**). Many genes significantly more accessible and expressed in progenitor dysfunctional cells in bulk RNA-seq analysis and bulk and single-cell ATAC-seq analysis (**Figure 1D, Figure 3I, Supplementary Figure 7**) were enriched in scRNA-seq cluster 9. Thus, cluster 9 consisted of progenitor and progenitor-like cells.

We next sought to characterize the expression profile of the progenitor cells and understand their relationship with the other cell states. Interestingly, cells in cluster 9 preferentially expressed genes induced during the early hours of T cell activation (“early response genes”), such as Xcl1, Cd200, Cxcl10, Cd69, Tnfsf8 (**Figure 4E,G, Supplementary Figure 13A**) (46–51). Xcl1 was the most strongly enriched gene in cluster 9; this gene was previously shown to be activated very early after antigen encounter and to be critical for engagement with antigen-presenting dendritic cells via Xcr1 (48, 49, 52); consistently, Xcr1+ dendritic cells were shown critical for promoting anti-tumor CD8 T cell immunity by anti-PD1 therapy via expansion of progenitor dysfunctional cells (53). On the other hand, many genes overexpressed in cluster 9 were also enriched in naïve cells, including Tcf7, Id3, Slamf6, Dapl1, Ccr7, Nsg2 (**Figure 4E,G**, **Supplementary Figure 13A**). This suggested that the progenitor and progenitor-like cells had a mixed gene expression profile resembling naïve, recently activated, dysfunctional, and memory cells.

Finally, we wanted to validate our observation from the scATAC-seq analysis of the similarity between progenitor-like cells in early chronic infection and their counterpart cells in the acute infection, likely memory precursor cells (**Figure 3E-H**, **Supplementary Figures 6E, 7A,B**). Indeed, scRNA-seq cluster 9 contained cells from all non-naïve samples (**Figure 4D**, **Supplementary Figure 13B**). Despite the limited resolution of differential expression analysis for these small subsets of cells, many of the critical marker genes were consistently overexpressed in this cluster when considered independently in d7 acute, d7 chronic, or d35 chronic infection, including many transcription factors and putative drivers of chromatin accessibility changes, such as Tox, Pou2f2, and Ikzf2; the most overexpressed gene in this cluster, Xcl1; the previously described progenitor markers Cxcr5 and Slamf6; and the memory precursor marker Il7r (**Figure 4G,H, Supplementary Figure 13C**). This again confirmed the emergence of progenitor-like cells early in chronic infection, together with a small number of similar cells in acute infection, that clustered with dysfunctional progenitor cells, consistent with recent reports (20, 54). We independently confirmed the presence of this CD8 T cell subpopulation at d7 in acute infection by reanalysis of three recently published scRNA-seq data sets (30, 31, 44) and confirmed enrichment of genes Tox, Ikzf2, Cxcr5 in this subpopulation **(Supplementary Figure 13D**). However, we also observed evidence of differential expression within the scRNA-seq cluster 9 between cells from different samples (**Supplementary Figure 14).** This again suggested that progenitor-like T cells at d7 in chronic infection may have already diverged from their analogs in acute infection, consistent with scATAC-seq analysis (**Figure 3G,J**, **Supplementary Figure 7C**).

Thus, our scRNA-seq analysis provided a comprehensive atlas of CD8 T cell functional and dysfunctional states and confirmed the existence of progenitor or progenitor-like cell population with shared features of naïve, memory and dysfunctional cells, and with highly common but still divergent transcriptional profiles between acute and chronic infection.

### scRNA-seq characterizes overlapping but divergent CD8 T cell responses to acute and chronic viral infection and cell-state specific transcription factor expression

We next sought to characterize the divergence between acute and chronic immune responses and expression of TF drivers using our scRNA-seq data set. Indeed, consistent with our scATAC-seq analysis (**Figure 3**), cells were already highly heterogeneous transcriptionally at d7 in both acute and chronic infection, occupying most of the expression landscape (**Figure 4I**). This included a small number of cells at d7 in acute infection that were terminally differentiated and similar to memory cells. Overall, cells in acute and chronic infection were markedly different as early as d7 (**Figure 4I**), again consistent with bulk chromatin state and gene expression analyses (**Figure 1B**, **Supplementary Figure 1F**) and with the scATAC-seq analysis (**Figure 3**). While cells in both infection settings occupied many of the same clusters, their proportions across clusters were different. Expression of certain genes also clearly differentiated between acute and chronic responses as early as d7. For example, the effector molecule Gzmb and cell division related genes Stmn1 and Rrm2 were overexpressed in acute infection, while the exhaustion marker Lag3, apoptosis gene Lgals1, and negative regulator of cytotoxicity and cell migration Pfn1 were overexpressed in chronic infection (**Supplementary Figure 11**). Thus, although acute infection predominantly drives CD8 T cell response along the effector cell trajectory, a small fraction of responding cells differentiate towards memory or even potentially along the dysfunctional trajectory. Conversely, chronic infection infrequently drives cells into an effector cell trajectory. This observation is consistent with previously reported findings (31). Together, our observations suggest that both chronic and acute infection can in principle give rise to cells at similar functional, activation, and differentiation states, but these cells accumulate at different proportions depending on the challenge.

Cell-state specific enrichment of TF expression (**Figure 4J**) was suggestive of cell state-specific function, consistent with observations from bulk and single-cell ATAC-seq data (**Figure 1D**, **Figure 2B,C**, **Figure 3G**). For example, we identified Gabpa as effector-specific, Rora as memory-specific, factors Tcf1, Nr4a1, Nfat5, Id3, Pou2f2, Pou6f1, Nfkb1, Batf as specific for progenitor dysfunctional state, and Zfp143, Zfp523 as specific for terminal dysfunction. Thus our scRNA-seq data supports the cell-state specific expression of TF drivers described in our bulk and single cell ATAC analyses.

### Allele-specific analysis validates cell state-specific transcription factor activity

To complement our observations of cell-state specific TF activities from the atlas of bulk and single-cell ATAC-seq and RNA-seq data in CD8 T cells across mouse models and immune challenges, we exploited our scRNA-seq profiling in F1 hybrid mice via allele-specific analysis of gene expression and TF binding, following and extending our previous study (43). Indeed, we identified thousands of genes with significant allele-specific expression in many clusters of our scRNA-seq data (**Supplementary Figure 15A**), including many genes involved in T cell activation and function (**Figure 5A**). Interestingly, most genes were consistently imbalanced in their expression towards B6 or Spret allele across clusters; however, we also identified a few genes, e.g. Gzmk or Fas, that were significantly imbalanced towards B6 or Spret in a cluster-dependent manner. We found that allelic imbalance of gene expression between B6 and Spret was correlated with differential expression between B6 and F1(B6xSpret) mice, confirming the accuracy of our gene expression allelic imbalance estimates (**Supplementary Figure 15B**).

**Figure 5.**
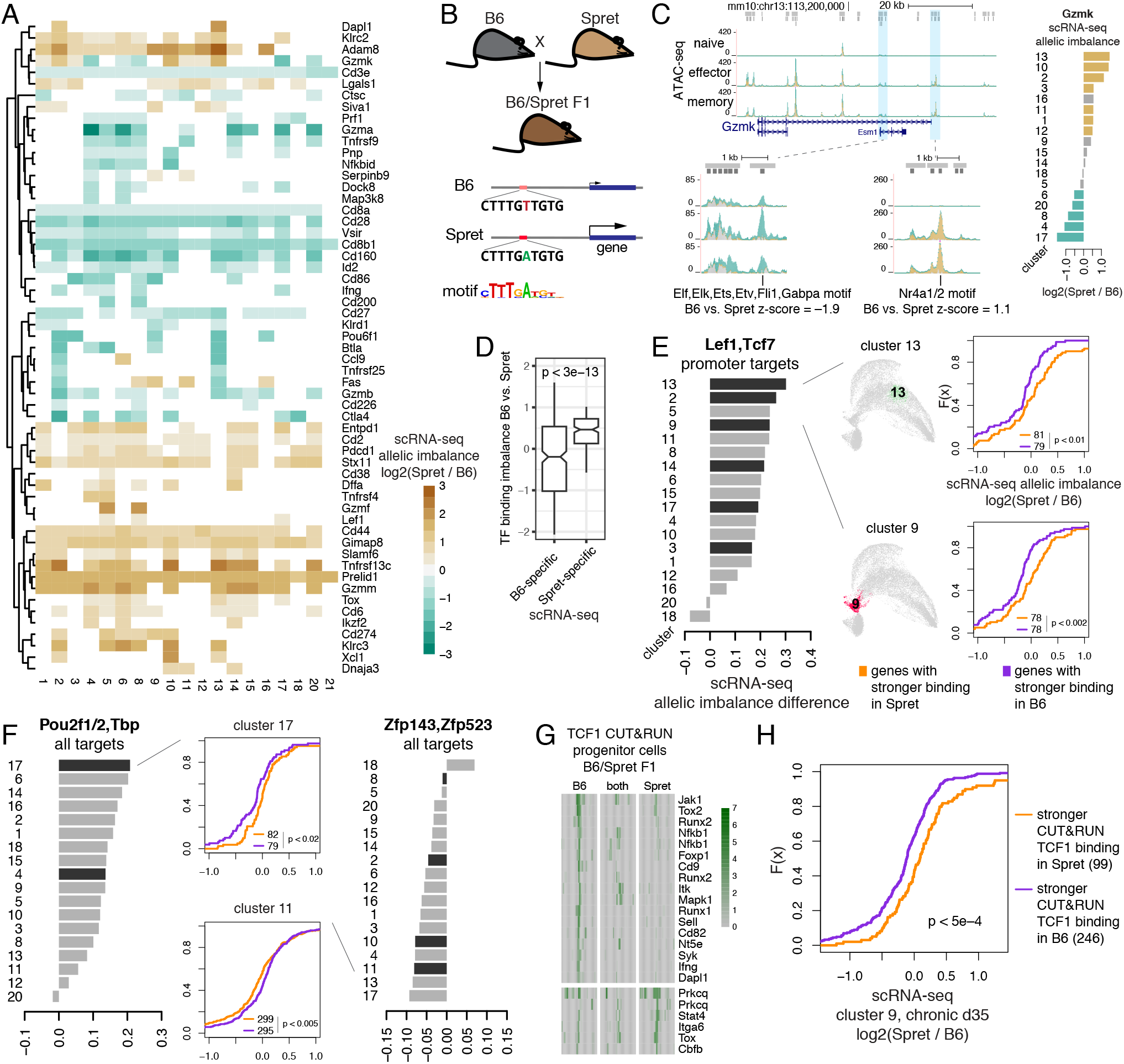
Allele-specific scRNA-seq analysis reveals cis-regulation of gene expression by transcription factors. **A.** Allelic imbalance of gene expression in clusters of scRNA-seq data. **B.** Schematic for analysis of association between predicted allelic imbalance of TF binding and allelic imbalance of gene expression. **C.** Left: ATAC-seq from B6/Spret F1 mice (43) at Gzmk locus and examples of peaks with B6-specific (green), Spret-specific (brown) and ambiguous (gray) ATAC-seq signal and allele-specific predicted TF binding. Right: allelic specificity of Gzmk expression with significant B6-specific (green) and Spret-specific (brown) expression highlighted. **D.** Predicted allelic imbalance of TF binding (difference between log odds scores of TF motif occurrence, z-score normalized) for genes with significant imbalance of expression towards B6 or Spret in any of the scRNA-seq clusters. Genes with only a single TF with predicted imbalanced binding are shown. **E-F.** Allelic specificity expression analysis of scRNA-seq data. CDF plots: allelic imbalance between B6 and Spret of library-size normalized scRNA-seq counts aggregated over cells in a selected cluster, for genes predicted to be bound more strongly in B6 or Spret using sequence motif analysis in 150bp windows around summits of their promoter peaks. Barplots: summary of the above analysis for each TF motif over all clusters; black bar indicates significant association (Kolmogorov-Smirnov p < 0.05). **G.** Examples of allele-specific TCF1 CUT&RUN binding sites in progenitor dysfunctional cells in the established chronic infection. **H.** Allelic imbalance between B6 and Spret of library-size normalized scRNA-seq counts aggregated over cells in cluster 9 at d35 in chronic infection, for genes bound by TCF1 more strongly in B6 or Spret as measured by CUT&RUN in progenitor dysfunctional cells.

To gain evidence for the causal role of TFs in regulating gene expression, we looked for an association between allele-specific TF binding, as predicted by motif analysis, and allele-specific expression (**Figure 5B,C**). For this, for each TF motif match in either B6 or Spret sequence around an ATAC-seq peak summit, we calculated TF motif binding imbalance as the difference in log-odds scores of motif match between B6 and Spret, z-score-normalized. Many genes, e.g. Gzmk (**Figure 5C**), had a lot of TFs with predicted binding imbalance at their promoter and enhancer ATAC-seq peaks. However, we expected that for genes with only a few potential TF regulators the relative effect of each regulator could be more impactful and therefore the association between allele-specific TF binding and allele-specific expression would be stronger. For example, when we restricted the analysis to genes with a single TF with predicted allele-specific binding, we found that the TF motif binding imbalance towards Spret allele was significantly higher for genes with significant expression imbalance towards Spret allele, and vice versa (**Figure 5D**). When focusing on genes with at most 20 ATAC-seq peaks, we found that imbalanced binding of certain TF motifs was significantly associated with differences in gene expression imbalance. For example, we found that genes with stronger TCF1/LEF1 promoter binding to Spret allele were significantly more imbalanced in their expression towards Spret than the genes with stronger binding of the same motif in B6 (**Figure 5E**). This suggests that TCF1/LEF1 motif binding in gene promoter is associated with activation of expression. Overall, using this approach, for multiple TFs we determined association of their binding with activation or repression of gene expression (**Figure 5F**, **Supplementary Figure 15C-E**).

TCF1 was previously reported as a critical marker of progenitor dysfunctional cells and is required for their generation during chronic infection (19, 21). To experimentally validate that TCF1 binding is associated with activation of gene expression, we mapped 3,325 TCF1 bound sites by CUT&RUN analysis in progenitor dysfunctional cells collected from F1(B6xSpret) hybrid mice at d35 upon infection with LCMV clone 13 (**Supplementary Figure 16A**). Direct TCF1 binding sites in the progenitor cells were strongly enriched for TCF1 motif (**Supplementary Figure 16B**) and included many T cell activation genes and markers of the progenitor state (**Supplementary Figure 16C,D**), including some targets with allele-specific TCF1 binding (**Figure 5G**, **Supplementary Figure 16E**). Consistent with previous observations and our predictions (**Figure 2**), accessibility levels at nearly all TCF1 targets were much higher in progenitor cells than in terminally dysfunctional cells (**Supplementary Figure 16F**). Surprisingly, these accessibility changes were not associated with changes in expression of nearby genes (**Supplementary Figure 16F**). However, allele-specific TCF1 binding was significantly associated with allele-specific expression of nearby genes in progenitor dysfunctional cells in the established chronic infection (**Figure 5H**), consistent with a causal role for TCF1 in activation of gene expression. Furthermore, allele-specific motif enrichment analysis in CUT&RUN-defined TCF1 binding sites revealed potential TCF1 binding co-factors, such as Runx and Nfkb family factors (**Supplementary Figure 16G**), consistent with allele-agnostic co-factor analysis (**Supplementary Figure 16B**).

These results demonstrate that allele-specific analyses in the hybrid genomes can reveal cis-regulatory effects of TF binding that may be missed with more traditional approaches.

### Progenitor cells differentiate to terminal dysfunction under acute viral challenge

Adoptive transfer experiments using sorted progenitor dysfunctional cells have demonstrated their potential to self-renew and give rise to terminally dysfunctional cells in models of melanoma and chronic viral infection (18, 20, 21). Since single cell chromatin and expression analyses showed that dysfunctional progenitors resemble progenitor-like cells at the plastic stage of T cell dysfunction development as well as the MPEC population in acute infection, we asked whether progenitor cells retain the potential to give rise to effector cells. We therefore transferred FACS sorted progenitor dysfunctional cells isolated from chronically infected mice on d35 post infection into congenically marked recipients infected with LCMV Arm (**Figure 6A,B**, **Methods**). The transferred cells were profiled at d7 post infection using flow cytometry and scRNA-seq. Flow cytometry revealed that the transferred CD8 T cells proliferated and persisted (**Figure 6C,D**). Furthermore, scRNA-seq analysis showed that these cells exhibited heterogeneous gene expression states (**Figure 6E-G**). Despite the acute infection setting, the phenotypic space defined by transferred cells after expansion most strongly overlapped with that of cells in chronic infection settings. Clustering analysis of the combined scRNA-seq data (**Supplementary Figure 17A,B**) confirmed that the expanded transferred cells were most similar in cluster composition to the cells at d7 in chronic infection (**Figure 6G**). Differential expression analysis between clusters limited to the expanded transferred cells also largely supported the cluster assignments of these cells (**Supplementary Figure 17C**), yielding largely the same differentially expressed genes as in the overall analysis (**Supplementary Figures 10,11**). Importantly, progenitor cells persisted upon transfer, confirming their self-renewal capacity (cluster t4, **Figure 6F**). These results suggest that the progenitor cells observed in chronic infection (d35) were present in acute infection settings and already committed to a dysfunctional fate.

**Figure 6.**
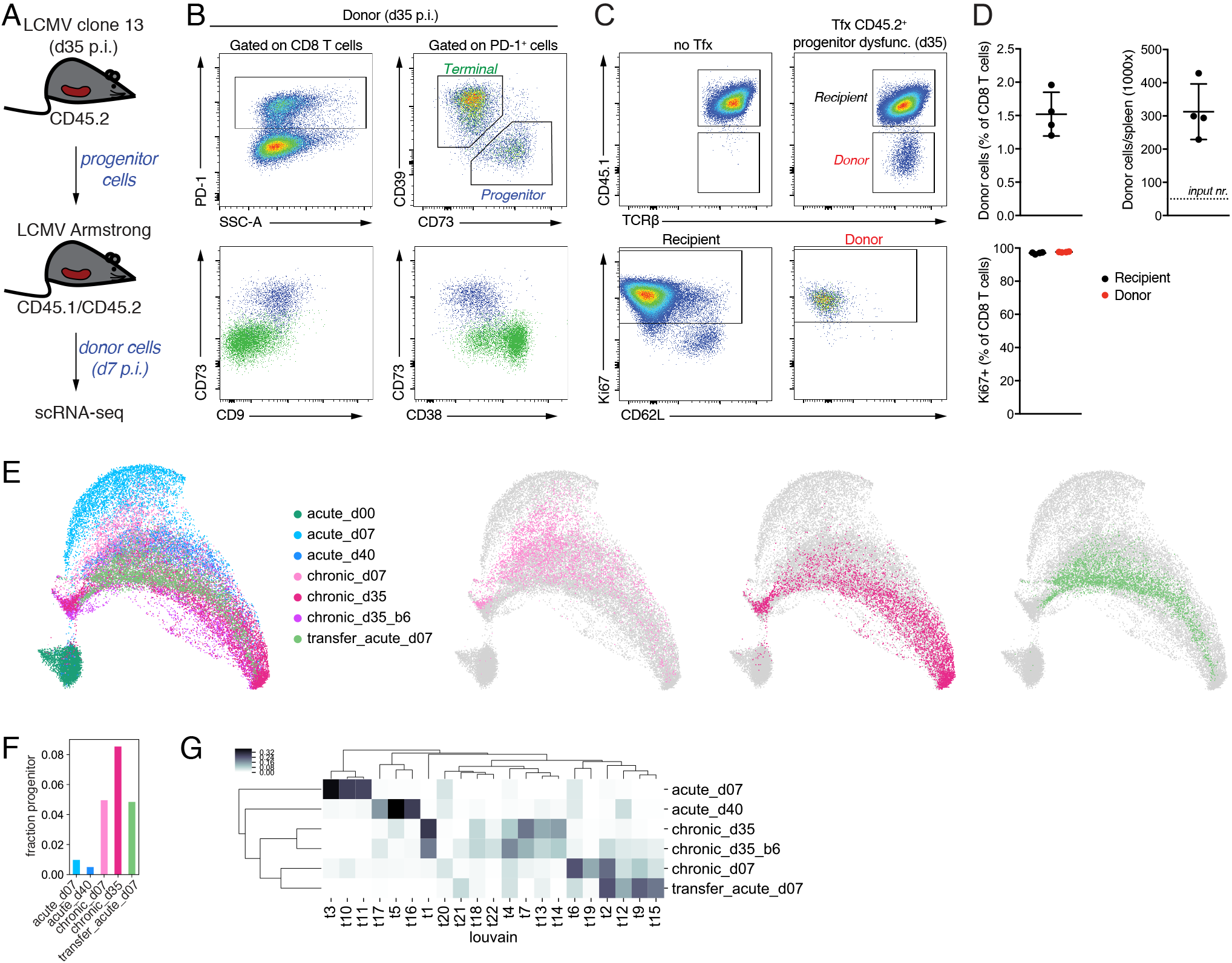
Progenitor CD8 T cells from chronic infection transferred to acute infection are committed to dysfunction. **A.** Setup for the adoptive cell transfer experiment. **B-C.** Flow cytometry of selected genes for progenitor cells before transfer (**B**) and after transfer and expansion (**C**) in acute infection. **D.** Quantification of flow cytometry results for donor cells after transfer and expansion. **E.** Left: UMAP of all scRNA-seq data including the transferred progenitor cells expanded in acute infection; right: UMAP showing individual samples within the overall map. **F.** Fraction of cells that belong to the progenitor cluster in individual samples. **G.** Cluster composition of samples. Heatmap showing for each cluster what fraction of cells (excluding naïve) that cluster occupies in each sample.

## DISCUSSION

Our unified analysis of bulk ATAC-seq and RNA-seq data across diverse studies of T cell response to chronic antigen exposure defined a universal program of progression to terminal exhaustion. By joint statistical analysis of data from numerous resources, we gained statistical power to robustly identify individual chromatin accessible sites, patterns of accessibility and expression changes at genic loci common across experimental models, and TFs whose binding motifs explain global accessibility patterns in distinct chromatin states in T cell differentiation.

Given the universality of T cell dysfunction program across mouse models, we then focused on one of the models, of LCMV infection, to elucidate the early divergence in chromatin states and transcriptional profiles between dysfunctional and functional T cell responses at single cell resolution using scATAC-seq and scRNA-seq. We found that a subset of antigen-specific T cells in chronic LCMV infection are appropriately activated and become effectors at day 7, while most progress along a dysfunctional trajectory. Conversely, a small number of cells in acute LCMV infection had progressed to a differentiated effector memory state, and potentially also to dysfunctional state by day 7. These findings are consistent with previous smaller-scale scRNA-seq studies of functional or dysfunctional T cells across tissues (18, 30, 31, 44, 45) but reveal a more nuanced comparison of acute vs. chronic immune responses, where different trajectories are possible but occur with different frequencies.

Critically, both in our scATAC-seq and scRNA-seq analyses, we found a TCF1+ progenitor-like cell population emerging as early as at day 7 post infection in both chronic and acute infection. In both infection settings, we found the same genes enriched in expression and chromatin accessibility at multiple promoter and enhancer peaks in this population, including Tcf7, Xcl1, Cxcl10, Ccr7, Slamf6, Id3, and Cxcr5, but also Tox and Ikzf2, which had previously been associated with terminally dysfunctional T cells. Notably, multiple lines of evidence from analyses of different data modalities converged on several TFs being strongly associated with this progenitor and progenitor-like population (**Figures 1D, 2B,C, 3G, 4J, 5E,F, Supplementary Figures 7B, 13A**), including the previously described TCF1 and the newly identified OCT2 (encoded by Pou2f2). However, differential accessibility analysis, motif regression modeling, and differential expression analysis identified subtle differences between cells from chronic and acute samples in this progenitor-like subpopulation, with cells from acute infection potentially representing a subset of IL7R^hi^ MPEC population and those from chronic infection forming a nascent progenitor reservoir. These findings may suggest an earlier divergence between progenitor and MPEC populations at the level of chromatin state and gene expression.

The presence of cells from both chronic and acute viral infection in the scRNA-seq cluster containing TCF1^+^ PD1^+^ progenitor cells and in the scATAC-seq progenitor-like cluster led us test the plasticity of the progenitor cell population. Using an adoptive cell transfer experiment together with scRNA-seq profiling and mapping these data to the comprehensive scRNA-seq atlas of CD8 T cell responses that we constructed, we confirmed that progenitor cells isolated from an established chronic viral infection were committed to a dysfunctional fate. While transferred progenitor cells could proliferate in a new host under acute immune challenge, they displayed the single cell phenotypic profile of a chronic immune response. These results are consistent with earlier adoptive T cell transfer studies showing that by day 15 post antigen encounter, T cells that had responded to chronic LCMV infection lost their ability to form functional memory cells under acute LCMV challenge (55) and with similar experiments documenting loss of plasticity for tumor-specific T cells in an autochthonous liver cancer model (56). However, these earlier studies also confirmed that at early time points in settings of chronic antigen stimulation, antigen-specific T cells – or at least a subpopulation of these cells – retained the potential to mount a functional response and form memory cells. Our single cell analysis of chromatin state and phenotypic heterogeneity of T cells under chronic immune challenge provides clues as to where this transient plasticity resides within the early T cell population. In particular, the similarity of the early progenitor-like population to the counterpart population in acute infection suggests that these cells may not yet be committed to progress to exhaustion, unlike TCF1^+^ PD1^+^ progenitor cells at later time points.

Our observations warrant further studies of the origins and regulatory mechanisms of maintenance and differentiation potential of the progenitor and progenitor-like cell populations across immune challenges, which could be performed in the future using parallel single cell TCR and transcriptomic profiling as well as heritable bar codes for tracking cell fates within a TCR clonal population over time (57) and at time points earlier than d7 upon T cell activation. Performing these studies and generating functional genomics data in F1 hybrid mice can further help interrogate regulatory mechanisms of gene expression by linking allele-specific analysis of TF motif occurrences (or allele-specific TF occupancy) to allelic imbalance of target gene expression.

In conclusion, our data sets and analyses lay the groundwork for resolving the fundamental questions of functional and dysfunctional T cell differentiation programs. We expect that our unified T cell atlas will provide a valuable resource both for basic T cell biologists and for cancer immunologists seeking to therapeutically target the transcriptional programs underlying progression to fixed dysfunction. Finally, our computational methods of analysis of bulk and single-cell ATAC-seq and RNA-seq data are broadly applicable.

## Acknowledgments

This study was supported by the NCI CSBC Research Center for Cancer Systems Immunology (U54 CA209975), the Alan and Sandra Gerry Metastasis and Tumor Ecosystems Center, and NCI Cancer Center Support Grant (P30 CA008748). Y.P. was supported by an AACR-Bristol-Myers Squibb Immuno-oncology Research Fellowship (19-40-15-PRIT) and Memorial Sloan Kettering-Parker Institute for Cancer Immunotherapy-Cycle for Survival Fund. We thank Lauren Fairchild for useful discussions and preliminary data analysis, Roshan Sharma and Manu Setty for advice on single-cell RNA-seq analysis, and members of the Leslie, Rudensky, and Pe’er labs for discussion and suggestions.

## SUPPLEMENTARY FIGURE LEGENDS

**Supplementary Figure 1. Universal program of CD8 T cell dysfunction (related to Figure 1). A.** Principal component analysis (PCA) of library-size normalized ATAC-seq read counts in 150bp windows around peak summits (functional cell state shown by color, data source by symbol) without batch-effect correction. **B.** Distributions of distances between ATAC-seq vectors of read counts in 150bp windows around peak summits. Distances were calculated in the two-dimensional PCA space built using 20000 most variable peak summit read counts, for pairs of samples in the same functional category (naïve, functional, dysfunctional), separately for pairs of samples from the same study and from different studies, before and after batch effect correction. **C.** PCA of library-size normalized batch-effect corrected ATAC-seq read counts in 150bp windows around peak summits (labels as in A) using 10000 most variable peak summit read counts. Here GLM-based batch effect correction used a factor encoding the functional cell state using two values, for naïve cells and all other cells. **D.** PCA of library-size normalized batch-effect corrected ATAC-seq read counts in 150bp windows around peak summits (labels as in A) using 10000 peak summit read counts most significantly differentially accessible between functional and dysfunctional cell samples. Here GLM-based batch effect correction used a factor encoding the functional cell state using two values, for naïve cells and all other cells; memory precursor cells and tissue-resident memory cells were not used in the differential accessibility analysis. **E.** Heatmap of accessibility for 10000 most significantly differentially accessible peaks between functional and dysfunctional cells. Library-size normalized ATAC-seq signal averaged over replicates is shown for 3000bp windows centered around peak summits. **F.** PCA of library-size normalized batch-effect corrected RNA-seq read counts for 1000 genes with most variable counts (functional cell state shown by color, data source by symbol). **G.** Scatter plot of differential expression (RNA-seq log2FC, x-axis) and differential accessibility (mean ATAC-seq log2FC over all peaks associated with a gene, y-axis) between functional and dysfunctional cells (computed as in D). Significantly differentially expressed genes shown as black dots. Significantly differentially accessible genes highlighted with red or blue color.

**Supplementary Figure 2. Differential accessibility and differential expression between functional and dysfunctional cells (related to Figure 1).**“Diamond” plots of differential accessibility and differential expression between functional and dysfunctional cells for genes associated with T cell activation, cytotoxicity, adhesion, and apoptosis, for transcription factors, for other cytokines and cytokine receptors, for other cell surface genes, and for other selected differentially accessible genes. In each panel, left: library-size normalized batch-effect corrected ATAC-seq read count log2FC for peak summits (diamond shown in color for significantly decreased/increased, FDR < 0.05) of significantly differentially accessible genes; right: log2 fold change of RNA-seq gene expression for the same genes.

**Supplementary Figure 3. Progression from early or progenitor to terminal dysfunctional state (related to Figure 1). A.** Comparisons of differential accessibility results. Each plot is a scatter plot of log2FC of library-size normalized batch-effect corrected ATAC-seq read counts in peak summit regions for two differential accessibility analyses. Color (blue, red, dark gray) is used to highlight quadrants with peak summit regions significantly differentially accessible in both pairwise comparisons. Counts in corners indicate the number of such peak summits in each quadrant. Spearman’s correlation is calculated over all such peak summits. **B.** Cumulative distribution function (CDF) of log-transformed quantile-normalized library-size normalized batch-effect corrected ATAC-seq read counts from various studies in 5940 peak summits significantly more accessible in progenitor than terminally dysfunctional T cells and in 2768 peak summits significantly more accessible in terminally than progenitor dysfunctional T cells in chronic LCMV infection (Kolmogorov-Smirnov p < 2e-16 for all comparisons). **C.** First principal component (PC1) of PCA for library-size normalized batch-effect corrected ATAC-seq read counts in dysfunctional T cells from different studies (see Figure 1C). PCA was calculated based on significantly differentially accessible peaks in different pairwise comparisons. For clarity, vertical random jiggle is added, and subsets of samples from different studies are shown below. **D.** Scatter plots of differential expression (RNA-seq log2FC, x-axis) and differential accessibility (mean ATAC-seq log2FC over all peaks associated with a gene, y-axis) between cell states. Significantly differentially expressed genes shown as black dots. Significantly differentially accessible genes highlighted with red or blue color. **E.** Cumulative distribution function of differential accessibility (ATAC-seq log2FC) between progenitor (TIM3-CD39-) and terminally (TIM3+ CD39+) dysfunctional TILs in human tumors (ATAC-seq data from Sade-Feldman *et al.* Cell 2018). Background distribution is for all peaks identified in human cells (black). Also shown are peaks evolutionarily conserved with peak summit regions in mouse melanoma model that were significantly more accessible in progenitor (blue) or terminally dysfunctional (red) cells; count and p-value from Kolmogorov-Smirnov test against the background distribution.

**Supplementary Figure 4. Differential accessibility and differential expression between early or progenitor and terminally dysfunctional cells (related to Figure 1).** Diamond plots of differential accessibility and differential expression (as in Supplementary Figure 2) between **A.** progenitor and terminally dysfunctional cells in chronic infection; **B.** progenitor and terminally dysfunctional cells in melanoma; **C.** dysfunctional cells profiled early (d8) and late (d27-28) in progression of chronic infection; **D.** dysfunctional cells profiled early (d5-7) and late (d14-60) in hepatocarcinoma progression.

**Supplementary Figure 5. Predictive models of transcription factor association with chromatin accessibility (related to Figure 2). A.** Schematic of the negative binomial generalized linear regression analysis to infer transcription factor (TF) associations with chromatin accessibility. **B.** Inferred TF motif coefficients for regressing library-size normalized batch-effect corrected ATAC-seq read counts in each sample, for motifs corresponding to coefficients with the highest variance across samples, z-score normalized within each row. **C.** Top: Boxplots of log2FC of RNA-seq gene expression between naïve and effector cells for genes associated with at least one peak significantly more accessible in naïve than effector cells and associated with at least one peak significantly more accessible in effector than in naïve cells. Left: default peak-gene associations (peak associated to the closest gene in genomic coordinates, if this gene is within 50Kbp). Center: peak–gene associations defined using significant Hi-C contacts in naïve cells. Right: peak–gene associations defined using non-significant Hi-C contacts in naïve cells. Bottom: same analysis for naïve vs. dysfunctional cells.

**Supplementary Figure 6. scATAC-seq analysis of CD8 T cells in acute and chronic infection (related to Figure 3). A.** Barplot showing the number of cells in each cluster from each sample. **B.** Components 2 and 3 of the UMAP representation of TF-IDF-transformed scATAC-seq data with Louvain clusters. **C.** Heatmap for single cells in scATAC-seq data, separately for acute and chronic infection, showing the naïve cell signature derived from bulk ATAC-seq data. **D.** Violin plots for scores in scATAC-seq data clusters, separately for acute and chronic infection, of peak signatures (for naïve cells, memory cells) derived from bulk ATAC-seq data. **E.** Violin plots for scores in scATAC-seq data clusters 1-8, 10, 13, separately for acute and chronic infection, of peak signatures (for effector cells, memory precursor cells, terminally dysfunctional cells) derived from bulk ATAC-seq data. **F.** Inferred TF motif coefficients for scATAC-seq counts averaged over cells in each cluster in each of the two samples. Inferred coefficients with the highest variance are shown (z-score row normalized).

**Supplementary Figure 7. scATAC-seq analysis reveals a similar population of precursor/progenitor cells in acute and chronic infection (related to Figure 3). A.** Genome browser tracks of ATAC-seq data for selected peaks. Bulk ATAC-seq for progenitor and terminally dysfunctional cells, and for terminal effector and memory precursor cells. Normalized aggregated single-cell ATAC-seq for cells in each cluster (for clusters 1-8, 10) in each of the two samples. For genes significantly more accessible in scATAC-seq cluster 10 as compared with other clusters. **B.** Each panel shows normalized scATAC-seq pseudocount (averaged over cells in each cluster in each sample) log2 fold change for peak summits of significantly differentially accessible genes; mean log2FC value is highlighted (with a shade of red) for genes that were overall differentially accessible. Shown are comparisons between cells in cluster 10 and cells in clusters 1-8, separately for acute and chronic infection, for genes highlighted in Figure 1E. **C.** Same as B, but for the most significantly differentially accessible genes between cells from acute and chronic infection within cluster 10.

**Supplementary Figure 8. Sorting for CD8 T cell subpopulations for scRNA-seq profiling (related to Figure 4). A-B.** Sorting strategy for isolating CD8 T cells responding to acute and chronic LCMV infection. **C.** Protein expression in cells isolated from chronic infection.

**Supplementary Figure 9. Dimensionality reduction and visualization of scRNA-seq data (related to Figure 4). A.** Principal component analysis (PCA) of library-size normalized batch-effect corrected ATAC-seq read counts in 150bp windows around peak summits (functional cell state shown by color, data source by symbol), including samples from naïve, effector, and memory cells from hybrid B6/Spret F1 mice. **B.** Projections of three-dimensional UMAP representation of library-size normalized scRNA-seq data. **C.** Log-transformed gene expression (library-size normalized scRNA-seq UMI counts) on UMAP. **D.** Cluster composition of samples. Heatmap showing for each cluster what fraction of cells that cluster occupies in each sample. **E.** Boxplot of the kNN graph distances from cells in samples “chronic_d35” (left) and “chronic_d35_b6” (right) to cells in other samples. For each cell C, average distance in the kNN graph from C to cells in each sample was calculated, and distribution of these values across cells C was plotted. **F-H.** Results of dimensionality reduction methods applied to normalized scRNA-seq data.

**Supplementary Figure 10. Differential expression analysis of scRNA-seq data (related to Figure 4). A.** Average scRNA-seq gene expression (library-size normalized UMI counts) in each cluster for differentially expressed genes between clusters (z-score row normalized). **B.** Barplots of scRNA-seq gene expression log2 fold change values for genes significantly differentially expressed in each cluster. **C.** MAGIC-imputed gene expression for selected genes. **D.** Fraction of cells classified as naïve (clusters 0, 7, 19) in individual samples.

**Supplementary Figure 11. Differential expression analysis within individual samples of scRNA-seq data (related to Figure 4).** Barplots of scRNA-seq gene expression log2 fold change values for genes significantly differentially expressed in each cluster in samples acute_d7 (**A**), acute_d40 (**B**), chronic_d7 (**C**), and chronic_d35 (**D**).

**Supplementary Figure 12. Association of bulk with single-cell RNA-seq data (related to Figure 4).** ssGSEA association of library size-normalized batch effect-corrected bulk RNA-seq data with library size-normalized scRNA-seq data averaged over cells in each cluster (z-score column normalized). Clusters 0, 7, 19 were most strongly associated with naïve cells, clusters 2, 10, 13 with memory cells, clusters 1, 3, 11, 12, 18, 21 with late and terminally dysfunctional cells, clusters 4, 5, 6, 8, 14, 15, 17, 20 with effector and early and progenitor dysfunctional cells, clusters 9 and 16 with progenitor cells and with early dysfunctional cells.

**Supplementary Figure 13. scRNA-seq cluster 9 consists of precursor/progenitor cells in acute and chronic infection (related to Figure 4). A.** Average scRNA-seq gene expression (library-size normalized UMI counts) in each cluster for genes significantly overexpressed in cluster 9 as compared with all other cells and as compared with all other cells except naïve cells (clusters 0, 7, 19). **B.** Fraction of cells that belong to cluster 9 in individual samples. **C.** MAGIC-imputed gene expression in individual samples. **D.** Enrichment of progenitor markers in a cluster of cells profiled at d7 in acute LCMV infection in three studies (30, 31, 44). Shown are significant (q < 0.01) log2 fold change values of expression in a selected cluster as compared with cells from all other clusters, as identified by diffxpy. Clusters occupying at least 1% of the total data set are labeled on UMAP.

**Supplementary Figure 14. Clustering and differential expression analysis for cells within scRNA-seq cluster 9 (related to Figure 4). A.** UMAP representation and more refined clustering of cells in scRNA-seq cluster 9 from samples acute_d7, acute_d40, chronic_d7, chronic_d35. Top: color by sample. Bottom: color by newly defined clusters p0–p4. **B.** Sample composition of clusters p0–p4. **C.** Average scRNA-seq gene expression (library-size normalized UMI counts) in clusters p0–p4 for selected differentially expressed genes between clusters (z-score row normalized). **D.** Barplots of scRNA-seq gene expression log2 fold change values for genes significantly differentially expressed in each of clusters p0–p4. These differential expression results suggest that cluster p3 consists of actively proliferating cells, clusters p0 and p1 are dominated by more mature progenitor dysfunctional cells observed in the established chronic infection, and clusters p2 and p4 are dominated by progenitor-like cells from d7 in acute or chronic infection. **E.** Barplots of scRNA-seq gene expression log2 fold change values for genes significantly differentially expressed between cells from sample acute_d7 that belong to cluster p4 and cells from sample chronic_d7 that belong to cluster p2.

**Supplementary Figure 15. Allele-specific analysis of TF binding and scRNA-seq data (related to Figure 5). A.** Allelic imbalance of gene expression in clusters of scRNA-seq data. **B.** Boxplots of differential expression (log2 fold change) between cells within the same scRNA-seq cluster profiled at d35 in chronic infection in B6 and in B6/Spret F1 mice, for genes with significant allelic imbalance towards B6 or Spret in B6/Spret F1 mice. Cluster number indicated on top of each boxplot. Significant difference between log2FC distributions (p < 2e–4 for clusters 2, 5, 10, 18, p < 2e–16 for all other clusters, Mann-Whitney U test) in all cases. **C-E.** Allelic specificity expression analysis of scRNA-seq data. CDF plots: allelic imbalance between B6 and Spret of library-size normalized scRNA-seq counts aggregated over cells in a selected cluster, for genes predicted to be bound more strongly in B6 or Spret using sequence motif analysis in 150bp windows around summits of their promoter peaks. Barplots: summary of the above analysis for each TF motif over all clusters; black bar indicates significant association (Kolmogorov-Smirnov p < 0.05).

**Supplementary Figure 16. CUT&RUN in progenitor dysfunctional CD8 T cells from B6/Spret F1 hybrid mice maps allele-specific TCF1 binding (related to Figure 5). A.** Normalized TCF1 CUT&RUN signal in TCF1-bound ATAC-seq peak summits (library-size normalized read counts in 50bp bins in 3000bp window around peak summit, ordered by total signal). **B.** Log2 fold change enrichment of motif frequency in TCF1 CUT&RUN peaks relative to all ATAC-seq peak summits. **C.** Examples of TCF1 binding sites. **D.** Genome browser tracks of bulk ATAC-seq for progenitor and terminally dysfunctional cells and TCF1 CUT&RUN in progenitor dysfunctional cells for selected loci. **E.** CDF curves for allelic imbalance of CUT&RUN signal between B6 and Spret for all TCF1 targets (black) and for those predicted to be bound more strongly in B6 (purple) or Spret (orange) using sequence motif analysis. **F.** Top: CDF plots of library size-normalized batch effect-corrected ATAC-seq signal for TCF1-bound sites vs. all ATAC-seq peak summits. Bottom: CDF plots of library-size normalized batch-effect corrected RNA-seq gene expression for genes with TCF1 binding vs. all expressed genes. Shown is the comparison between progenitor and terminally dysfunctional cells in chronic LCMV infection. **G.** Imbalance of predicted TF binding using sequence motif analysis in peaks with stronger TCF1 binding in B6 or in Spret allele as identified by CUT&RUN.

**Supplementary Figure 17. Differential expression analysis of the transferred and expanded cells profiled with scRNA-seq (related to Figure 6). A.** Clustering of scRNA-seq data, including the sample of transferred progenitor cells that expanded in acute infection, using the Louvain algorithm. **B.** Comparison of new clustering with the previous one (shown in Figure 4C). Shown is a fraction of cells in each of the clusters t0–t23 that belong to each of the previously obtained clusters 0–21. This analysis suggests that cluster t4 almost coincides with previously obtained cluster 9 and therefore consists of progenitor and progenitor-like cells. **C.** Barplots of scRNA-seq gene expression log2 fold change values for genes significantly differentially expressed in cells from each cluster in sample transfer_acute_d7.

## METHODS

### Bulk ATAC-seq and RNA-seq analysis

#### Construction of ATAC-seq peak atlas

ATAC-seq data from multiple previous studies were downloaded from GEO using fastq-dump from NCBI SRA Toolkit (https://trace.ncbi.nlm.nih.gov/Traces/sra/sra.cgi?view=toolkit_doc). Functional annotation of samples (naïve, effector, memory, dysfunctional, etc.) was obtained from each study. Reads were aligned to the mouse genome mm10.GRCm38 using bowtie2 v2.3.4.3 (58). Uniquely aligned reads were extracted using SAMtools v1.9 (59). Peaks were called using MACS2 v2.1.1.20160309 (60). For each data source, peaks were called using all samples from all replicates combined. Then IDR was used to identify reproducible peaks between at least some pair of replicates of the same condition. The union of all such peaks formed peak library for each study. Then peaks identified in each study were all combined into a common list, merging overlapping peaks from different studies, and filtered based on ATAC-seq read coverage. This resulted in an atlas of 129799 chromatin accessibility peaks for CD8 T cells. Peaks were further split into subpeaks around summits of signal, using a custom script based on the signal processing package https://bitbucket.org/leslielab/biosignals. Peaks were annotated using GENCODE vM14 (61). Each peak was associated with the closest gene, if this gene was within 50Kb in genomic coordinates. Peaks were annotated by applying the following sequence of rules: peaks were classified as promoter peaks if within 2Kb from a transcription start site of any annotated transcript; otherwise as exonic if overlapping with any exon of any annotated transcript; otherwise as intronic if within a gene body of any annotated gene; or finally as intergenic if within 50Kb of a gene. All the remaining peaks were left unclassified. Reads from each sample were counted in peaks and 150bp genomic regions centered around peak summits using Rsubread v1.32.4 (62).

#### Batch effect correction for ATAC-seq

DESeq2 v1.22.2 (63) was used to fit multi-factorial models to ATAC-seq read counts in peaks or peak summits. The main model included two factors where one factor represented the source of the data, and the other factor represented the functional annotation of the sample, broadly defined as naïve, functional, and dysfunctional. For PCA analysis and visualization, batch effect-corrected values were used. To do the batch effect correction, log2FC values associated with the data source factor were extracted from the model for all peaks and used to correct the original counts, by dividing them by the exponentiated log2FC values.

The alternative variant of the model included a two-valued functional annotation factor, where one value was for naïve samples, and the other for all antigen-experienced cells, including all functional and dysfunctional cells. This alternative variant of the model produced results similar to the main model (Supplementary Figure 1C).

In order to determine whether memory precursor cell samples and tissue-resident memory cell samples were similar to functional or dysfunctional cells, a variant of this alternative model with a two-valued functional annotation factor was used. For this, differentially accessible peaks were detected between functional and dysfunctional cells but excluding memory precursor and tissue-resident memory cell samples. Then PCA analysis for all samples was run on ATAC-seq read counts restricted to these differentially accessible peaks, revealing that memory precursor and tissue-resident memory cells are more similar to other functional effector and memory cells (Supplementary Figure 1D).

#### Differential accessibility analysis

DESeq2 v1.22.2 was used to perform differential accessibility analysis between selected pairs of conditions. For this, original counts were used, and the model included factors for functional annotation, cell state, data source, and/or type of immune challenge, where appropriate.

#### Transcription factor motif analysis

Transcription factor (TF) binding motifs for *Mus musculus* were downloaded from CIS-BP version 1.02 (64) via the web interface (compressed archive Mus_musculus_2016_06_01_2-46_pm.zip). For the DNA sequences in 150bp-wide regions around peak summits, script findMotifsGenome.pl from HOMER suite (65) was run with parameters ‘‘mm10 -len 8,10,12 -size given -S 100 -N 1000000 -bits -p 10 - cache 1000’’ in order to identify the significance of presence of each motif in the sequences of the peaks as compared with the background sequences. We limited the analysis to motifs corresponding to expressed TFs, defined as those with at least 200 library-size normalized RNA-seq reads in at least one condition in at least one study. We focused only on the motifs present in at most 50% of the peak summit region sequences. Furthermore, we detected *de novo* motifs using HOMER in the same sequences and associated them with TFs by similarity with the CIS-BP motifs using script compareMotifs.pl with parameters “-reduceThresh 0.7 -matchThresh 0.9”. The most significant motif per TF, either from the database or identified *de novo*, was selected for further analysis (with potentially multiple TFs associated with the same motif), if it had HOMER p-value < 0.001. This resulted in 113 motifs. Furthermore, we merged motifs that had correlation of motif occurrences in the peak summits above 0.75. This resulted in the list of 105 motifs corresponding to 204 TFs for further analysis. FIMO version 4.11.2 (66) was used to search for motif matches in 150bp regions around the peak summits. Matches with p-value < 1e–3 were chosen as significant.

#### Predictive modeling of transcription factor activity

To infer cell-state specific TF activities, we performed a supervised modeling of chromatin accessibility data based on TF motif occurrences. We formed a feature matrix that consisted of TF motif match predictions in regions around peak summits. Each value in the matrix was a sum of FIMO log-odds scores of a TF motif occurrence in a peak summit region. For each ATAC-seq sample, we performed negative binomial generalized linear regression modeling of the batch-effect corrected chromatin accessibility values ****y**** in peak summit regions using this feature matrix, with ridge regularization, using function cv.glmreg() from R package mpath (67) with parameters family = “negbin”, alpha = 0, theta = 1, nfolds = 5, maxit = 20000, thresh = 1e-5. We limited the regression analysis to peak summit regions with at least 10 batch-effect corrected reads on average across all samples. We identified a hyperparameter multiplier ****a**** of the ridge regularization penalty term using 5-fold cross-validation; for this we ran the regression with 30 values of ****a**** formed by multiplying the mean of ****y**** by a vector of values from 10-8 to 10^3^ (with equidistant log_10_ values) and choosing the value of ****a**** that maximizes the Spearman correlation between the observed and predicted values of ****y****. Then we used the coefficients of this regression as a proxy to TF activity scores and used them for downstream analysis; for this, we limited our downstream analysis to results of regression only for those ATAC-seq samples where the fit converged for at least 4 values of the hyperparameter ****a****.

#### RNA-seq data analysis

RNA-seq data from multiple previous studies were downloaded from GEO using fastq-dump. Functional annotation of samples (naïve, effector, memory, dysfunctional, etc.) was obtained from each study. Reads were aligned to the mouse genome mm10.GRCm38 using hisat2 v2.1.0 (68). Reads from each sample were counted in genes annotated by GENCODE vM14 using Rsubread v1.32.4. Batch effect correction and differential expression analysis were performed the same as for ATAC-seq data.

#### Comparison of ATAC-seq and RNA-seq using Hi-C contacts

Published Hi-C data from naïve CD8 T cells (39) was used to define associations of ATAC-seq peaks to genes and compare it with our default peak-to-gene association (see *Construction of ATAC-seq peak atlas*), using a computational experiment comparing ATAC-seq and RNA-seq data. Hi-C read pairs were processed using HiC-Pro pipeline (version 2.11.1) (69). Reads were trimmed to 30nt and mapped to the mm10 reference genome. We used valid pairs output in cis from HiC-Pro that were mapped uniquely to the genome. Valid pairs from replicates were merged and binned into 10kb bins. HiC-DC (70) was used to assign statistical significance to each interaction bin. Significant chromatin loops between 10Kb bins were defined at q-value < 0.05 and with support by more than 20 Hi-C valid pairs, resulting in 22,337 significant loops. Median distance between loop ends was 180Kb, and only 5% of loops had distance between ends shorter than 50Kb. For control, non-significant loops were defined as those with q-value between 0.1 and 0.2 and supported by not more than 10 Hi-C valid pairs, resulting in 118,497 loops. Peaks were associated to genes using these loops as follows. For a loop between 10Kb-long genomic regions A and B, a peak was associated to a gene if the peak was within 5Kb window from A and another peak at a promoter of the gene was within 5Kb from B. Then differential gene expression (RNA-seq log2FC) between naïve and effector cells was compared between genes associated with at least one significantly more accessible peak in naïve than in effector cells and in effector than in naïve cells (FDR < 0.05), for different methods of association of peaks to genes (Supplementary Figure 5C): default associations, associations based on significant loops, and associations based on non-significant loops. The same analysis was performed for comparison between naïve and dysfunctional cells (Supplementary Figure 5C).

### Single-cell ATAC-seq experiments

Four male C57BL/6J mice per group were infected with either LCMV Armstrong (2×10^5^ p.f.u. via intraperitoneal injection) or LCMV Clone 13 (2×10^6^ p.f.u. via retroorbital injection). On day 7 post-infection, total CD8 T cells from pooled spleens were double sorted by flow cytometry and prepared for single cell ATAC-seq analysis.

scATAC-seq was performed using Chromium instrument (10x Genomics) and Single Cell ATAC Reagent Kits (Chemistry v1). The suspension of cells was processed following the User Guide (CG000168 Rev A). Briefly, cells were lysed in bulk, washed and resulting nuclei suspension treated with ATAC reagents provided in the kit. Approximately, 8000 nuclei per sample were loaded onto microfluidics Chip E and encapsulated with DNA barcodes and reaction mix. Following emulsion-PCR the resulting material was purified and subjected to 12-cycles of indexing PCR. The indexed scATAC-Seq libraries were double-size selected using SPRI beads and sequenced on an Illumina NovaSeq 6000 instrument (Read 1 – 50 cycles, i7 index – 8 cycles, i5 index – 16 cycles, and Read 2 – 50 cycles) at 125M reads per sample.

### Single-cell ATAC-seq data analysis

#### Preprocessing, dimensionality reduction, clustering

The data was preprocessed with cellranger-atac from 10X. Then using the BAM files with read alignments for the two samples produced by cellranger-atac, peaks were detected using MACS2. An extended list of peaks was formed by combining the previously constructed bulk ATAC-seq peaks with the non-overlapping newly identified peaks. The cellranger-atac tool was then rerun with this extended peak list as input to obtain a count matrix for each of the two samples. Cells with a log library size less than 3.5 and peaks that were accessible in less than 4 cells were excluded from the analysis. The count matrix was then binarized and transformed using term frequency-inverse document frequency (TF-IDF, scikit-learn package v0.20.3 (https://scikit-learn.org/) for normalization, followed by PCA and UMAP for dimensionality reduction and visualization. Cells were clustered by the Louvain clustering method. Signature peak sets for progenitor, naïve, effector, MPEC, and terminal dysfunction were derived from bulk ATAC-seq analysis and then used to score cells by taking the average normalized counts of the peak set and subtracting the average normalized counts of a reference peak set. Analysis and visualizations were performed using python package scanpy v1.4.4 (71).

#### Comparison with bulk ATAC-seq data compendium

BAM files were generated with aligned scATAC-seq reads corresponding to cells from each cluster, further split between the two samples. Pseudobulk counts were obtained from these BAM files for the extended peak list using Rsubread v1.32.4. Limited only to clusters 1-6, 8, 10, 13 that contained activated virus-specific cells, differential accessibility analysis was then run using these pseudobulk counts for each of the clusters, within each of the two samples, vs. all the remaining clusters split between the two samples. The most significantly differentially open peaks formed a signature set of peaks for each cluster split between the two samples. Then ssGSEA (40) was run for these signatures against batch-effect corrected library-size normalized bulk ATAC-seq counts using the package GSVA v1.30.0 (72).

### Single-cell RNA-seq experiments

#### Isolation of cells for scRNA-seq

For isolation of CD8 T cells responding to acute infection, B6/Spret F1 mice, where B6 stands for C57BL/6J and Spret stands for SPRET/EiJ, were infected with 2×10^5^ p.f.u. of LCMV Armstrong via intraperitoneal injection or left uninfected. Splenocytes were stained with a cocktail of fluorescent antibodies, NP396 tetramer, and viability dye to mark dead cells, and sorted using flow cytometry. Naïve CD8 T cells were isolated from pooled spleens of 2 uninfected mice as CD44-CD62L+ CD8 T cells. Activated and memory effector cells were isolated as NP396+ CD8 T cells from pooled splenocytes of 2 or 3 mice on day 7 or day 40 post-infection, respectively and used as input for scRNA-seq analysis.

For isolation of cells responding to chronic infection, B6/Spret F1 or B6 mice were infected with LCMV clone 13 by retroorbital injection of 4×10^6^ p.f.u. of virus. Mice were depleted of CD4 T cells by injection of 200ug aCD4 antibody (BioXCell cat: BE0003-1) one day prior to and one day post-infection. Cells were isolated as live CD62L-CD8 T cells from spleens of 3 pooled mice per group and used as input for scRNA-seq analysis.

#### Adoptive transfer of progenitor dysfunctional cells

4 C57BL/6J mice were infected with LCMV Clone 13 (2×10^6^ p.f.u. via retroorbital injection). On day 35 post-infection, splenocytes were pooled and stained using a cocktail of fluorescently labeled antibodies and Ghostdye Violet 510 (Tonbo cat: 13-0870-T100) to distinguish dead cells. Dysfunctional progenitor cells were isolated by flow cytometry-based sorting as PD1+, CD73+, CD39-CD8 T cells. This population was also CD9+ and CD38 low. 50,000 dysfunctional precursor cells/recipient were transferred into 4 CD45.1/CD45.2 mice via retroorbital injection. Recipients were infected with LCMV Armstrong (2×10^5^ p.f.u. via intraperitoneal injection). On day 7 post-infection donor cells were re-isolated based on expression of congenic markers and prepared for scRNA-seq.

#### Single-cell barcoding, library preparation and sequencing

Single-cell suspensions were loaded on 10x Genomics Chromium instrument and encapsulated in microfluidic droplets with barcoded DNA hydrogel beads and RT reagents from Chromium Single Cell 3’ Reagent kit (v3). The cDNA synthesis/barcoding was performed following manufacturer’s instructions: 53°C for 45 min followed by heat inactivation at 85°C for 5 min. The barcoded-cDNA was purified and PCR-amplified and prepared for sequencing according to the Single Cell 3’Reagent kit 3 User Guide (CG000183; Rev B). The DNA sequencing was performed on the Illumina NovaSeq 6000 instrument (R1 read – 26 cycles, R2 read – 70 cycles or higher, and index read – 8 cycles), aiming for ∼200 million reads per ~5,000 single cells.

### Single-cell RNA-seq data analysis

#### Preprocessing, dimensionality reduction, clustering

We analyzed scRNA-seq data from the six samples labeled “acute_d00”, “acute_d07”, “acute_d60”, “chronic_d07”, “chronic_d35”, “chronic_d35_b6”. Reads were aligned to the combined B6 and Spret genome using hisat2 v2.1.0. For this, files Mus_musculus.GRCm38.dna.toplevel.fa.gz and Mus_spretus_spreteij.SPRET_EiJ_v1.dna.toplevel.fa.gz with genomic sequences of B6 and Spret mice, respectively, were obtained from NCBI FTP server, and hisat2 index for the chromosomes from both files was constructed. Using the BAM files of scRNA-seq read alignment, UMI counts for each gene in B6 and Spret genomes were obtained using a custom script, by overlapping read alignments with exonic annotations of genes in either B6 or Spret and counting UMI corrected using the method from seqc (73). For this, files Mus_musculus.GRCm38.91.gtf.gz and Mus_spretus_spreteij.SPRET_EiJ_v1.86.gtf.gz with gene annotations for B6 and Spret genomes, respectively, were obtained from NCBI FTP server. The resulting read count tables were used for downstream analysis in scanpy v1.3.7. Cells with less than 100 genes with positive counts or cells with more than 40000 total positive UMI counts were filtered out, and genes with less than 500 positive counts across cells from all samples were filtered out. Cells with high expression of B cell genes (likely contamination) were filtered out (15 cells). Ribosomal genes were excluded from the analysis, and only protein-coding genes were included in the analysis. Based on inspection of the distribution of the number of genes detected per cell in each sample, cells from sample “acute_d00” with log_10_ gene count less than 2.85, from sample “acute_d07” with log_10_ gene count less than 2.9, from sample “acute_d60” with log_10_ gene count less than 2.9, from sample “chronic_d07” with log_10_ gene count less than 2.9, from sample “chronic_d35” with log_10_ gene count less than 2.8, from sample “chronic_d35_b6” with log_10_ gene count less than 2.8 were excluded from the analysis. Then the count matrix was filtered again to include only cells with at least 500 genes and genes with at least 100 counts. This resulted in a read count table of 9822 genes in 24400 cells across the six samples.

Counts for 9124 genes present in both B6 and Spret annotations were used for normalization, dimensionality reduction, and clustering. For normalization, the total count in each cell was calculated, excluding the top 50 genes with the highest total count across all cells, and then each count divided by the total count per cell and multiplied by the median of total counts per cell. For dimensionality reduction and visualization, principle component analysis (PCA) was run for the normalized counts, the first 50 principle components (PCs) were selected, and the kNN (nearest neighbor) graph was built for k = 50 nearest neighbors per cell using Euclidian metric. The kNN graph was then used to construct three-dimensional uniform manifold approximation and projection (UMAP) with default parameters. The data was also visualized with t-distributed stochastic neighbor embedding (tSNE) using function “TSNE” from package sklearn.manifold with perplexity = 150 and otherwise default parameters applied to the first 50 PCs of normalized count matrix; with diffusion maps and force-directed atlas applied to the kNN graph. Louvain clustering was then applied to the kNN graph with resolution = 1.9 and otherwise default parameters. For visualization of gene expression, imputation algorithm MAGIC (74) was applied to the normalized count matrix using package magic with parameters a=15, k=30, knn_dist=’euclidean’, n_pca=50, random_state=0, t=3.

To assess the similarity of samples “chronic_d35” and “chronic_d35_b6”, kNN graph analysis was performed. For each cell c and a sample S, the distance between c and S was calculated as the average distance in the kNN graph between c and cells from S. This value was calculated for each cell c from samples “chronic_d35” and “chronic_d35_b6” against each of the samples, and shown as a boxplot (Supplementary Figure 9E).

#### Differential expression analysis

Differential expression analysis between cells from groups A and B was performed as follows. Let x_G_ and y_G_ be vectors of normalized scRNA-seq expression values of the gene G in individual cells from groups A and B, respectively. Then log2 fold change (log2FC) of expression for G is estimated as log_2_((Y_G_ + c) / (X_G_ + c)) where X_G_ = mean(x_G_), Y_G_ = mean(y_G_) are means of normalized expression of G in cells from groups A and B, respectively, and c is a corrective value defined as median over all values M_g_ where M_g_ is the mean expression of a gene g across all cells from all samples. Only genes with the absolute log2FC above a certain threshold were then tested for significance. For each such a gene G, the significance of the log2FC for G was estimated with the Mann-Whitney U test applied to normalized counts x_G_ and y_G_ for G in individual cells from groups A and B; the Bonferroni correction for multiple hypothesis testing was then applied to these p-values.

Using this approach, differential expression analysis was performed for each cluster vs. cells from all other clusters, testing for significance of all absolute log2FC values above 0.5. The same analysis was also repeated when excluding all naïve cells defined as cells from sample “acute_d00” or clusters 0, 7, 19. Differential expression analysis was also performed for each pairwise comparison between clusters. Furthermore, for each sample, differential expression analysis was performed for each cluster vs. cells from all other clusters when restricting the analysis only to cells from that sample.

#### Comparison with bulk RNA-seq data compendium

The most significantly differentially expressed genes in each scRNA-seq cluster (adjusted p < 0.001, log2FC > 0.8), defined as the union of at most 100 genes from the comparison of the cluster with all other clusters and at most 100 genes from the same comparison restricted to non-naïve cells, formed a signature set of genes for this cluster. ssGSEA was then run for these signatures against batch-effect corrected library-size normalized bulk RNA-seq read counts using package GSVA v1.30.0.

#### Analysis of scRNA-seq data from adoptive cell transfer experiment

The six main scRNA-seq samples were combined with the sample “transfer_acute_d07” from the progenitor dysfunctional cells extracted from the established chronic infection and adoptively transferred and expanded under acute infection. The extended data set was preprocessed in the same manner as the main data set, resulting in a count matrix for 9274 genes in 28447 cells, including 4047 cells from the transfer sample. The analysis of this extended count matrix was performed in the same manner as with the main six samples, resulting in clusters t0–t23. This cell clustering, restricted to the cells from the main six samples, was compared with the previously obtained clusters, suggesting cluster t4 consisted of the progenitor and progenitor-like cells. The cluster composition of each sample, calculated as fraction of cells in each sample that belongs to each cluster, was also compared between samples, suggesting the sample “transfer_acute_d07” was most similar to the sample “chronic_d07”.

### Allelic specificity analysis

#### Allele-specific scRNA-seq expression

For allele-specific analysis, scRNA-seq sequencing reads were re-aligned in allele-specific manner as previously described (43, 75). Briefly, the genetic variants of Spret mice were obtained from the mouse genome project (76). A pseudo-Spret genome was built by modifying the reference genome with SNPs, insertions and deletions found in the wild-derived inbred strain. The RNA-seq reads were aligned to the reference and pseudo Spret genomes in parallel using STAR (77) with the following parameter setting: with the following parameter settings: $STAR --runMode alignReads --readFilesCommand zcat -- outSAMtype BAM --outBAMcompression 6 --outFilterMultimapNmax 1 --outFilterMatchNmin 30 -- alignIntronMin 20 --alignIntronMax 20000 --alignEndsType Local. Next, the genomic coordinates of pseudo-genome aligned reads were converted back to the corresponding B6 coordinates. To determine the allelic origins of the reads, the mapping scores of the two alignments of each reads were compared. We retained the alignment with the highest score and generated the final BAM files. In cases where the diploid genome alignment produced identical scores for both genomes, one of the alignments was selected randomly.

Spret- and B6-specific and ambiguous UMI counts of gene expression for each cell were obtained with custom scripts, in the same manner as in the main analysis (see *Single-cell RNA-seq data analysis)*. Further analysis was restricted only to cells and genes selected for the main scRNA-seq analysis. Library-size normalization for Spret- and B6-specific and ambiguous read counts was obtained applying the same scaling for each cell as in the main analysis. The allelic imbalance of expression of each gene in each scRNA-seq cluster was defined as log2((w_1_ + w_2_ / 2) / (w_0_ + w_2_ / 2)), where w_0_ and w_1_ are B6- and Spret-specific total library-size normalized counts for this gene in this cluster, respectively, and w_2_ is the ambiguous count for this gene in this cluster. Significance of the allelic imbalance was assessed by a Mann-Whitney U test applied to all B6-specific vs. all Spret-specific library-size normalized gene expression estimates for this gene over cells in this cluster.

#### Allele-specific predicted TF binding

Spret sequences of bulk ATAC-seq peak summit regions, as defined above (see *Construction of ATAC-seq peak atlas)*, were obtained by introducing genetic variants between B6 and Spret into B6 sequences of these regions. To estimate allelic specificity of predicted TF binding, FIMO was run for Spret sequences and rerun for B6 sequences of the peak summit regions with the same motif collection as described previously (see *Transcription factor motif analysis)* with a relaxed p-value threshold of 5e-3. All matches at p < 1e-4 in either B6 or Spret sequence were selected, and allelic imbalance of predicted TF binding for each motif in each peak summit region was estimated as the difference between the total FIMO log odds score for all matches of this motif in Spret and in B6. Then the distribution of these values for this motif across all peaks was z-score-normalized.

Analysis of the association of allelic imbalance of predicted TF binding and of scRNA-seq gene expression was performed for each TF motif and each scRNA-seq cluster as follows. All predicted TF motif binding sites with substantially stronger predicted match at Spret or B6 sequence were selected as those with z-score above 0.2 or below −0.2, respectively. Then the scRNA-seq allelic imbalance in this cluster was compared between genes nearby B6-specific and Spret-specific predicted binding sites, with significance assessed by Kolmogorov-Smirnov test.

### CUT&RUN experiments

Terminal and progenitor dysfunctional cells were isolated from the spleens of 4 male B6/Spret F1 mice or B6 mice on day 35 post-infection, based on expression of cell surface markers (see *Adoptive transfer of progenitor dysfunctional cells*). Cells from 2 mice were pooled to generate biological duplicates with approximately 70,000 cells per replicate. CUT&RUN libraries were prepared as described (78) with the modifications described below. Because Concanavalin-A (ConA) is a well-known T cell mitogen, we avoided the use of ConA-coated beads for cell isolation and handling. 70k cells per replicate were collected in a V-bottom 96 well plate by centrifugation and washed in antibody buffer (buffer 1 (1x permeabilization buffer from eBioscience Foxp3/Transcription Factor Staining Buffer Set diluted in nuclease free water, 1X EDTA-free protease inhibitors, 0.5mM spermidine) containing 2mM EDTA). Cells were incubated with TCF1 antibody for 1h on ice. After 2 washes in buffer 1, cells were incubated with pA/G-MNase at 1:4000 dilution in buffer 1 for 1h at 4 degrees. Cells were washed twice in buffer 2 (0.05% (w/v) saponin, 1X EDTA-free protease inhibitors, 0.5mM spermidine in PBS) and resuspended in calcium buffer (buffer 2 containing 2mM CaCl2) to activate MNase. Following a 30 minute incubation on ice, 2x stop solution (20mM EDTA, 4mM EGTA in buffer 2) was added and cells were incubated for 10 minutes in a 37 degree incubator to release cleaved chromatin fragments. Supernatants were collected by centrifugation and DNA was extracted using a Qiagen MinElute kit.

CUT&RUN libraries were prepared using the Kapa Hyper Prep Kit (Kapa Biosystems KK8504) and Kapa UDI Adapter Kit (Kapa Biosystems KK8727) according to manufacturers protocol with the modifications described below. A-tailing temperature was reduced to 50 degrees to avoid melting of short DNA fragments and reaction time was increased to 1h to compensate for reduced enzyme activity as described by Liu *et al.* 2018. Following the adapter ligation step, 3 consecutive rounds of Ampure purification were performed using a 1.4x bead to sample ratio to remove excess unligated adapters while retaining short adapter-ligated fragments. Libraries were amplified for an average of 15 cycles using a 10 second 60 °C annealing/extension step to enrich for shorter library fragments. Following amplification, libraries were purified using 3 consecutive rounds of Ampure purification with a 1.2x bead to sample ratio to remove amplified primer dimers while retaining short library fragments. A 0.5x Ampure purification step was included to remove large fragments prior to sequencing.

### CUT&RUN data analysis

#### Identification of transcription factor binding sites

CUT&RUN reads from B6 mouse samples were aligned to the genome using bowtie2 v2.3.4.3. Similar to RNA-seq, paired-end CUT&RUN reads from F1 B6/Spret mouse samples were mapped to the diploid genome using STAR with the splicing alignment feature switched off. The command line was as follow: $STAR --runMode alignReads --genomeLoad NoSharedMemory --readFilesCommand zcat -- outSAMtype BAM SortedByCoordinate --outFilterMultimapNmax 1 --outFilterMatchNmin 40 -- outBAMcompression 6 --outFilterMatchNminOverLread 0.4 --seedSearchStartLmax 20 --alignIntronMax 1 --alignEndsType Local. We only used the read pairs if their fragment length is between 50 to 500 bp.

For analysis, we used two control IgG samples and biological replicates for TCF1 CUT&RUN, including two from B6 mice and two samples from F1 B6/Spret mice. CUT&RUN and control reads were counted in 150bp windows around ATAC-seq peak summits. To estimate library sizes, control regions were obtained by shifting ATAC-seq peak summit regions by 2Kb in either direction, extending the shifted segments by 500bp preserving their center, and excluding those that overlapped with ATAC-seq peak summit regions, and then calculating control and CUT&RUN read counts in these control regions and using DESeq2 v1.22.2, as described previously (79). Differential CUT&RUN count analysis was then run using DESeq2, identifying 3325 TCF1 binding sites in the progenitor dysfunctional cells defined as ATAC-seq peak summit regions with significantly higher TCF1 CUT&RUN read counts than IgG control (adjusted p-value < 0.1). To produce heatmaps of TCF1 CUT&RUN signal around binding sites, bigWig files and count matrices in bins around binding sites were produced using deepTools (80).

### Transcription factor motif enrichment analysis

Motif enrichment in TCF1 binding sites was estimated as log2FC of frequency of TCF1 binding sites with significant TCF1 motif match (FIMO p < 1e-4) over such frequency for all ATAC-seq peak summit regions, with significance of enrichment estimated using hypergeometic test followed by Bonferroni correction for multiple hypothesis testing.

### Allele-specific analysis

For the TCF1 binding sites, allelic imbalance of TCF1 binding was defined as log2((w_1_ + w_2_ / 2) / (w_0_ + w_2_ / 2)), where w_0_ and w_1_ are B6- and Spret-specific CUT&RUN counts, respectively, and w_2_ is the count of ambiguously aligned reads. For the CDF plots of allelic imbalance of TCF1 binding (Supplementary Figure 16E), TCF1 binding sites with predicted stronger TCF1 motif batch in Spret or B6 were selected as those with TCF1 motif match imbalance z-score above 0.1 or below −0.1, respectively. For the allele-specific TCF1 co-factor analysis, Spret- or B6-specific TCF1 binding sites were detected as those with log2FC of CUT&RUN read count in Spret over B6 above 0.5 or below −0.5, respectively. Then for these sets of allele-specific TCF1 binding sites, for each motif, co-factor motif enrichment was calculated by comparing motif match imbalance z-scores against those in all ATAC-seq peak summit regions, with significance estimated using hypergeometric test. Motif match imbalance z-scores were then plotted as a boxplot (Supplementary Figure 16G) for all the motifs that were detected as significant both for Spret- and B6-specific TCF1 CUT&RUN binding sites.

