## Supplementary Figures for "A unified atlas of CD8 T cell dysfunctional states in cancer and infection"

#### Supplementary Figure 1

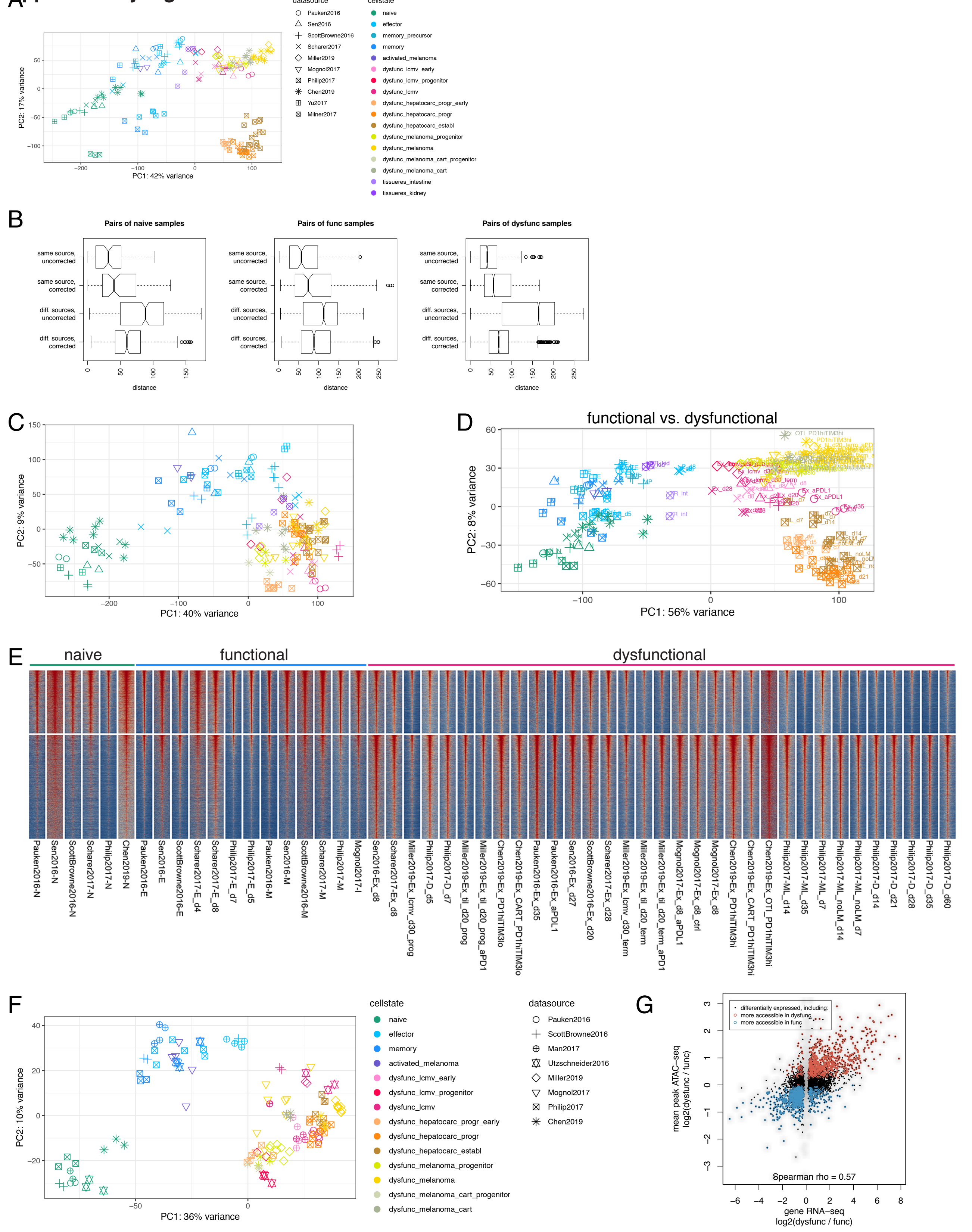

### Supplementary Figure 2

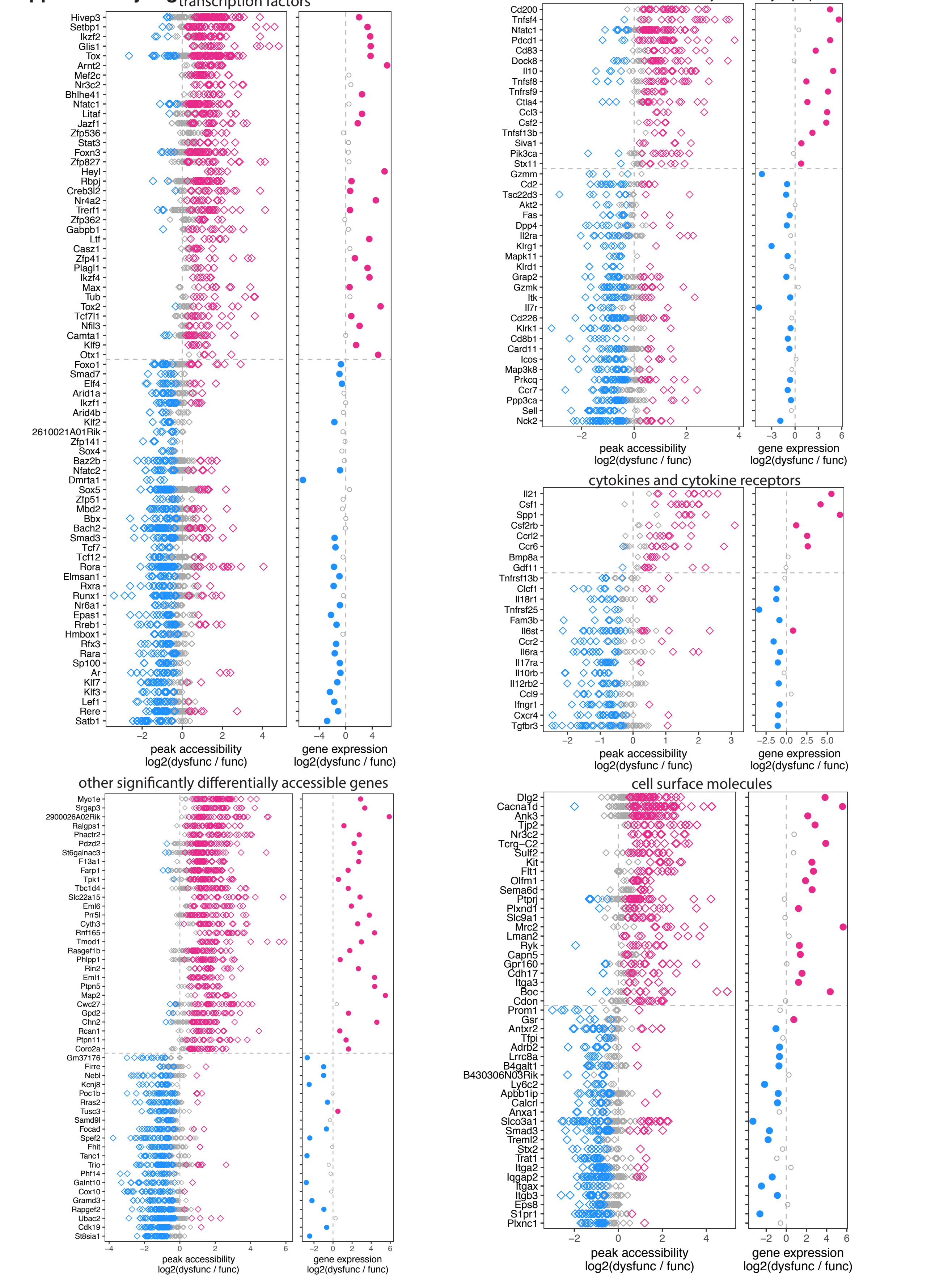

Supplementary Figure 3

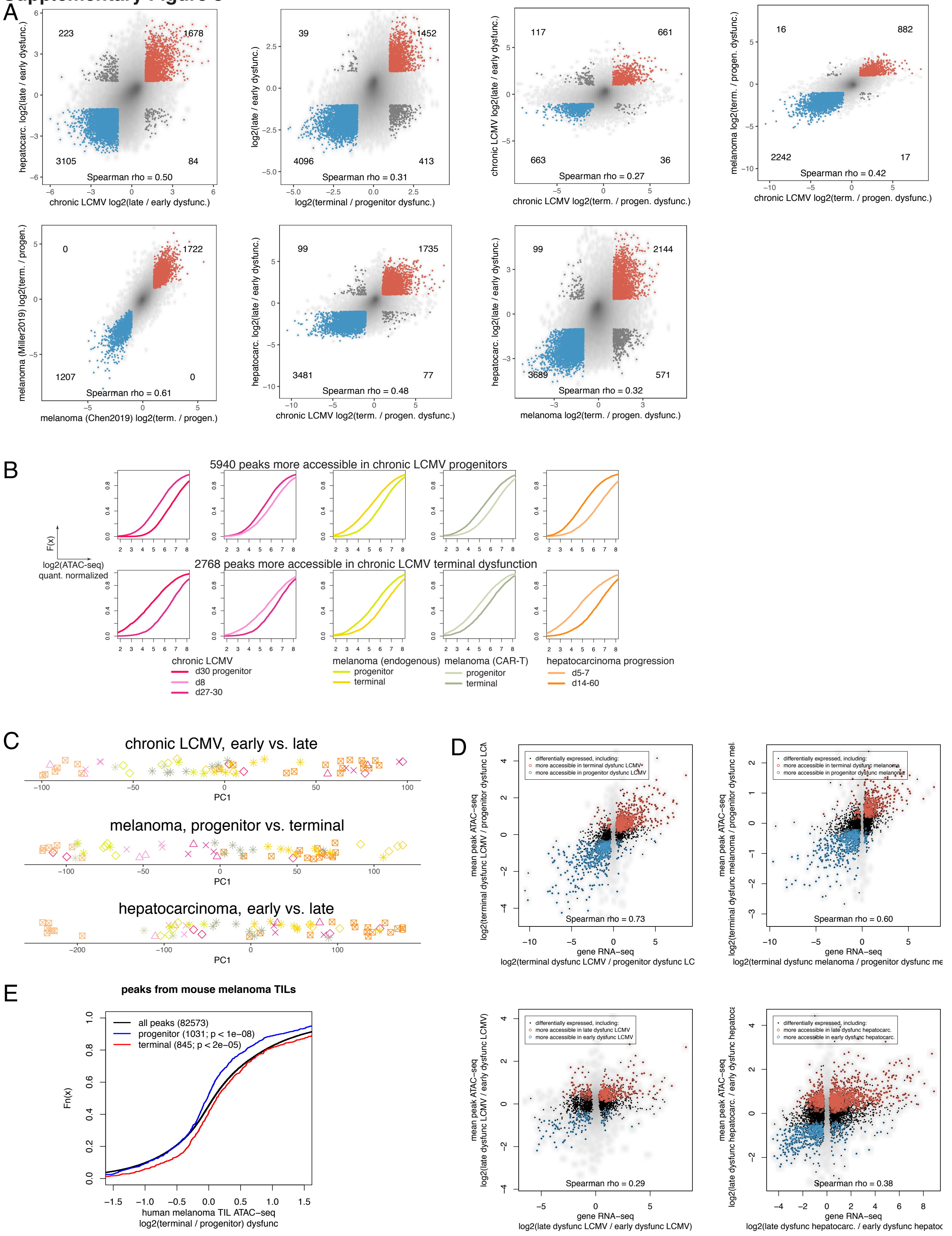

### Supplementary Figure 4

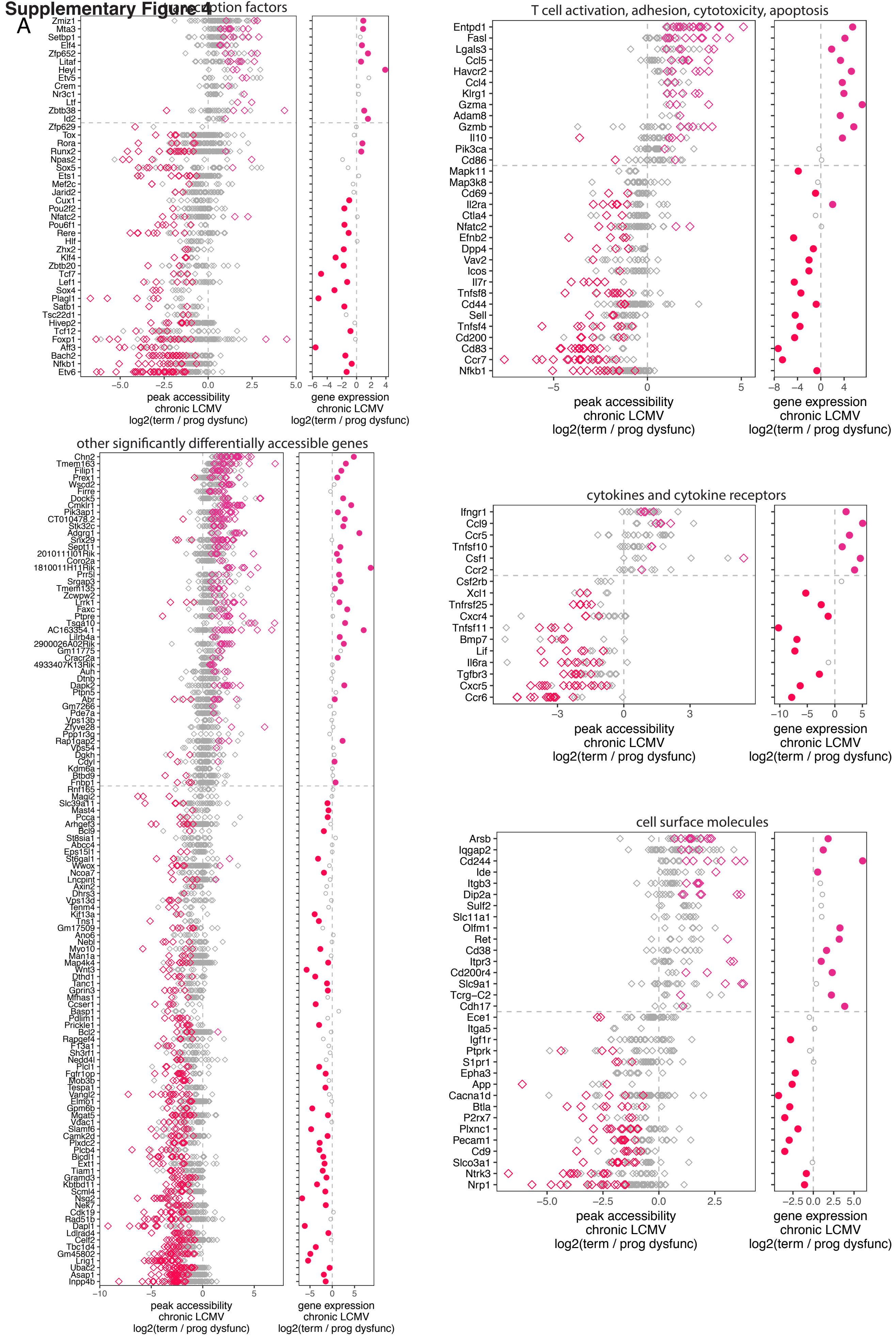

### Supplementary Figure 4

B

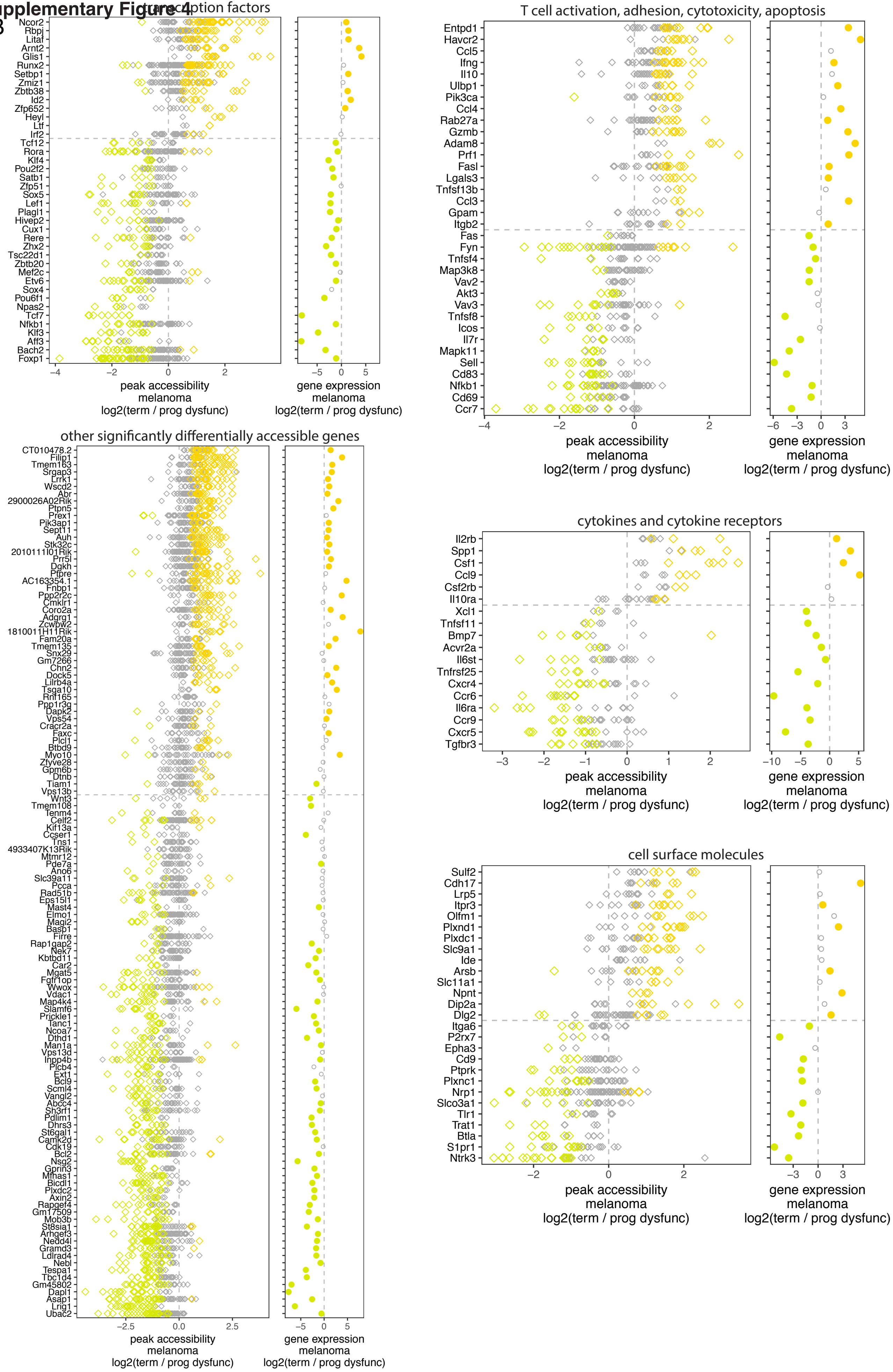

### Supplementary Figure 4

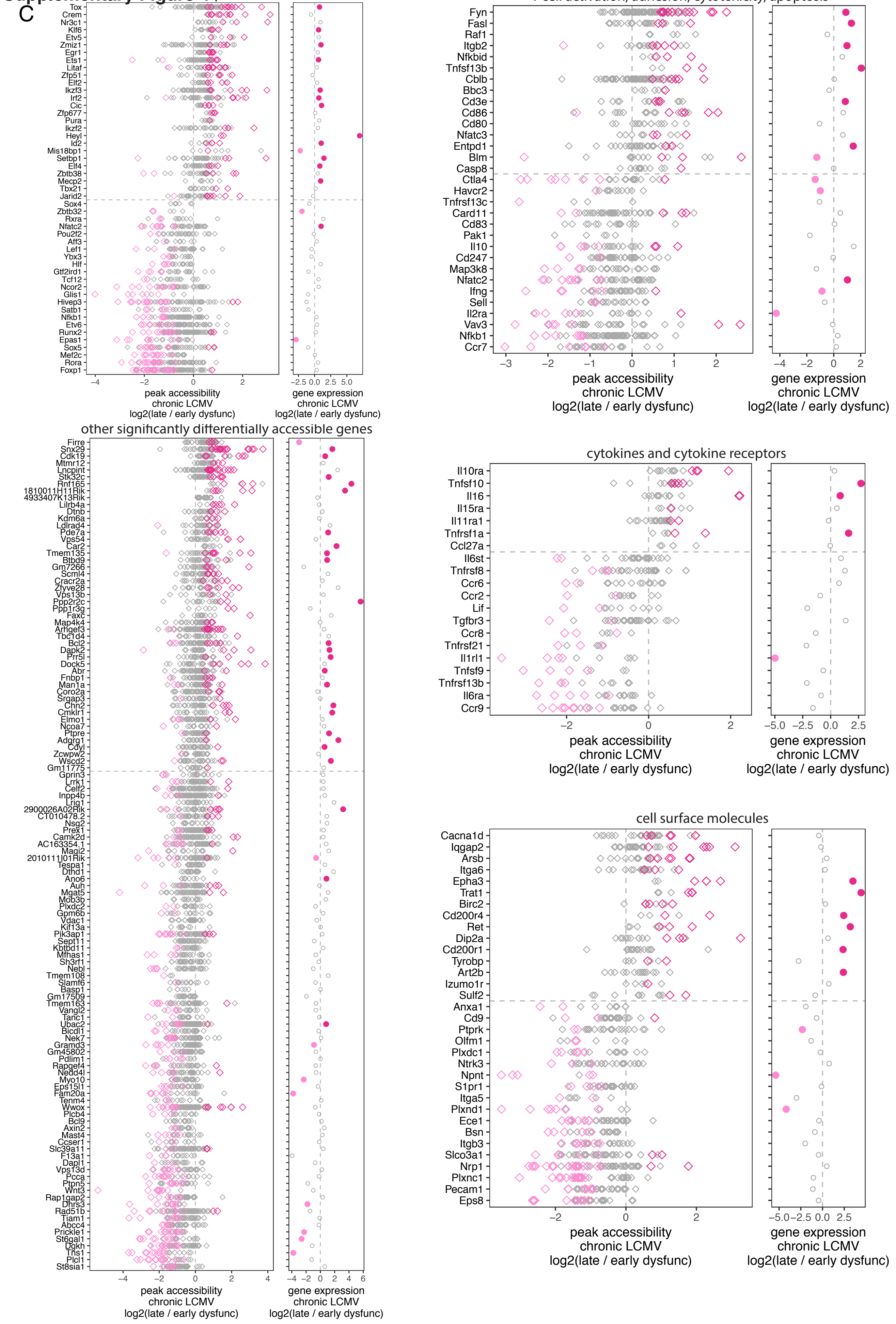

**Supplementary Figure 4**

D

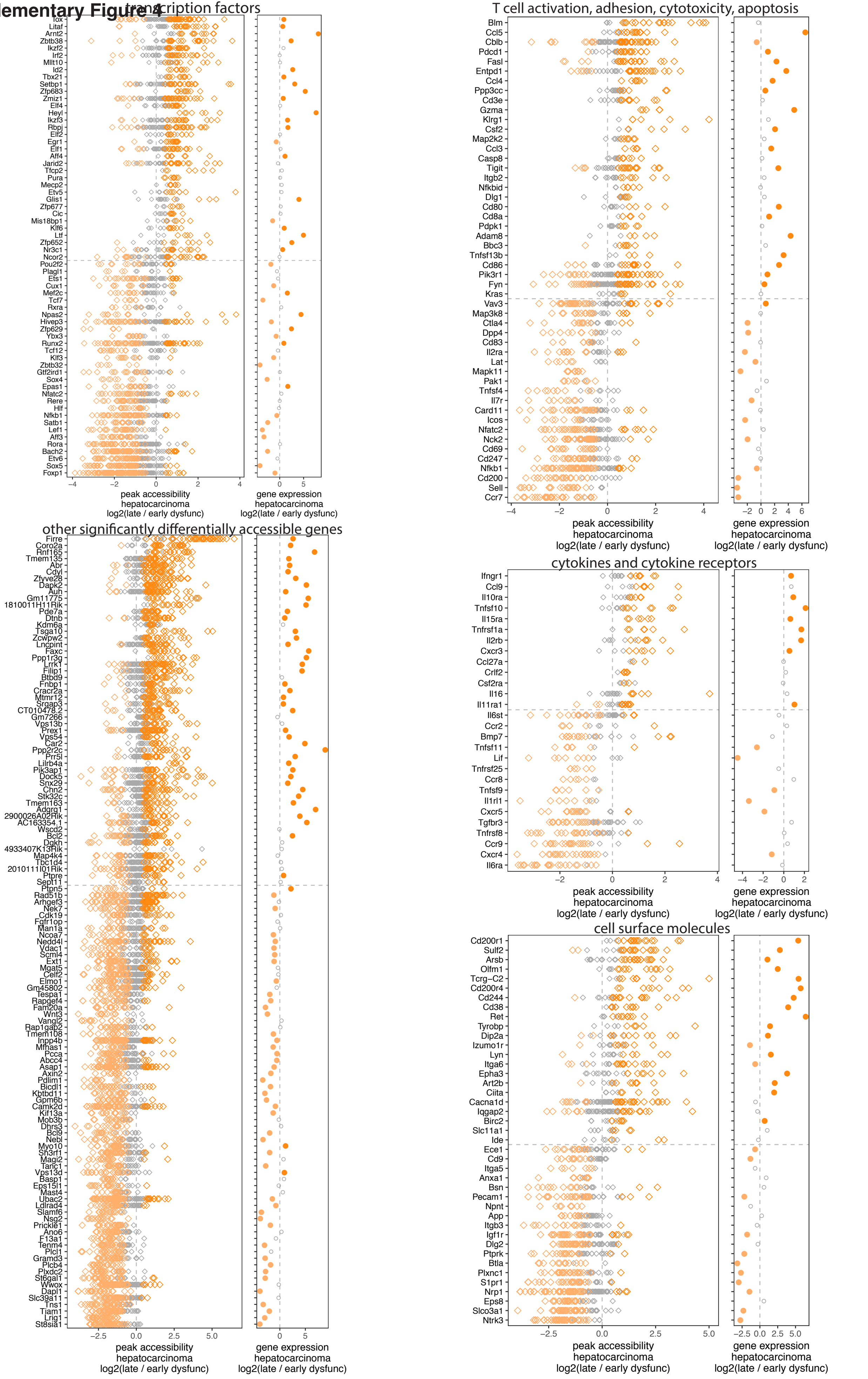

**Supplementary Figure 5**

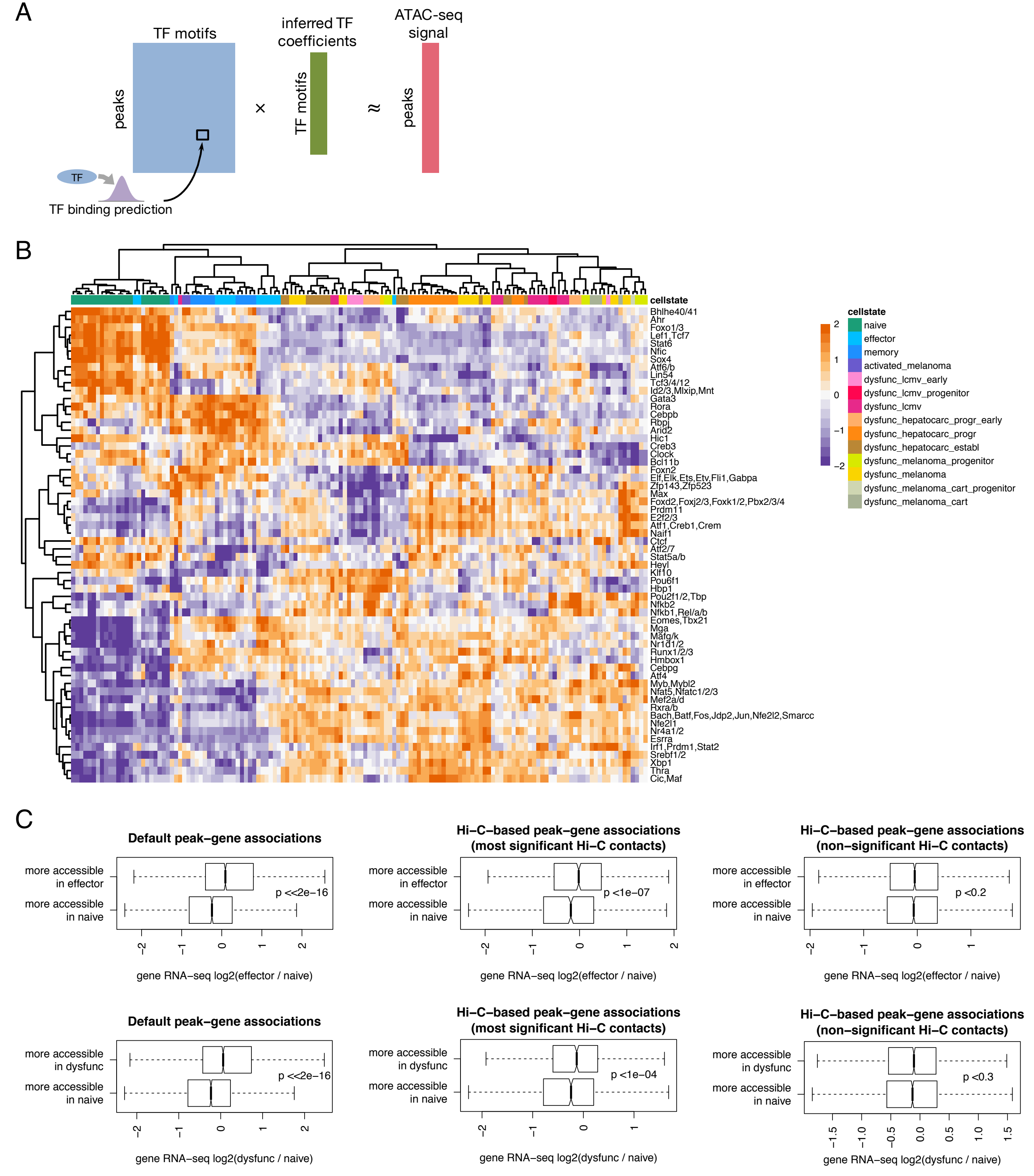

### Supplementary Figure 6

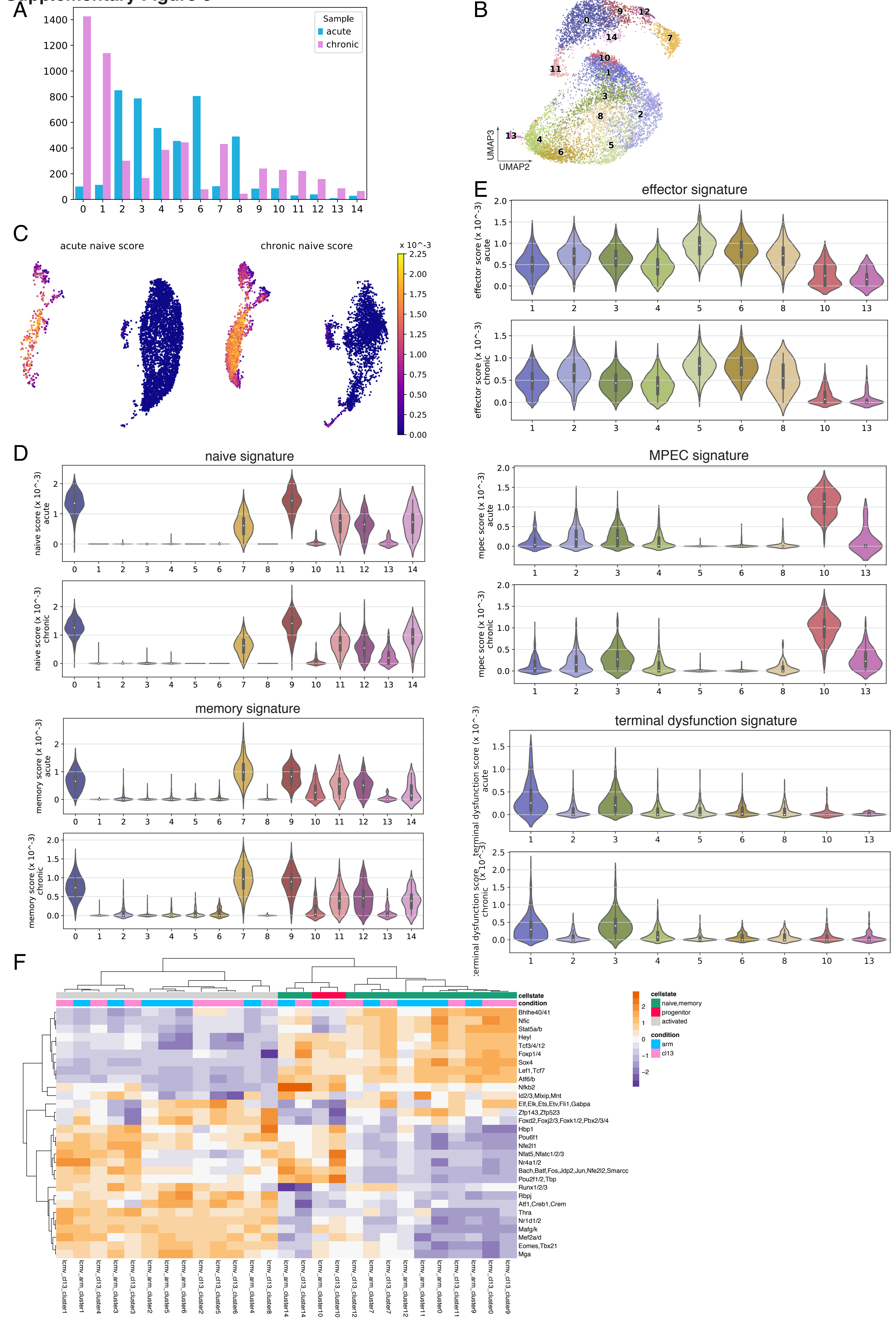

### Supplementary Figure 7

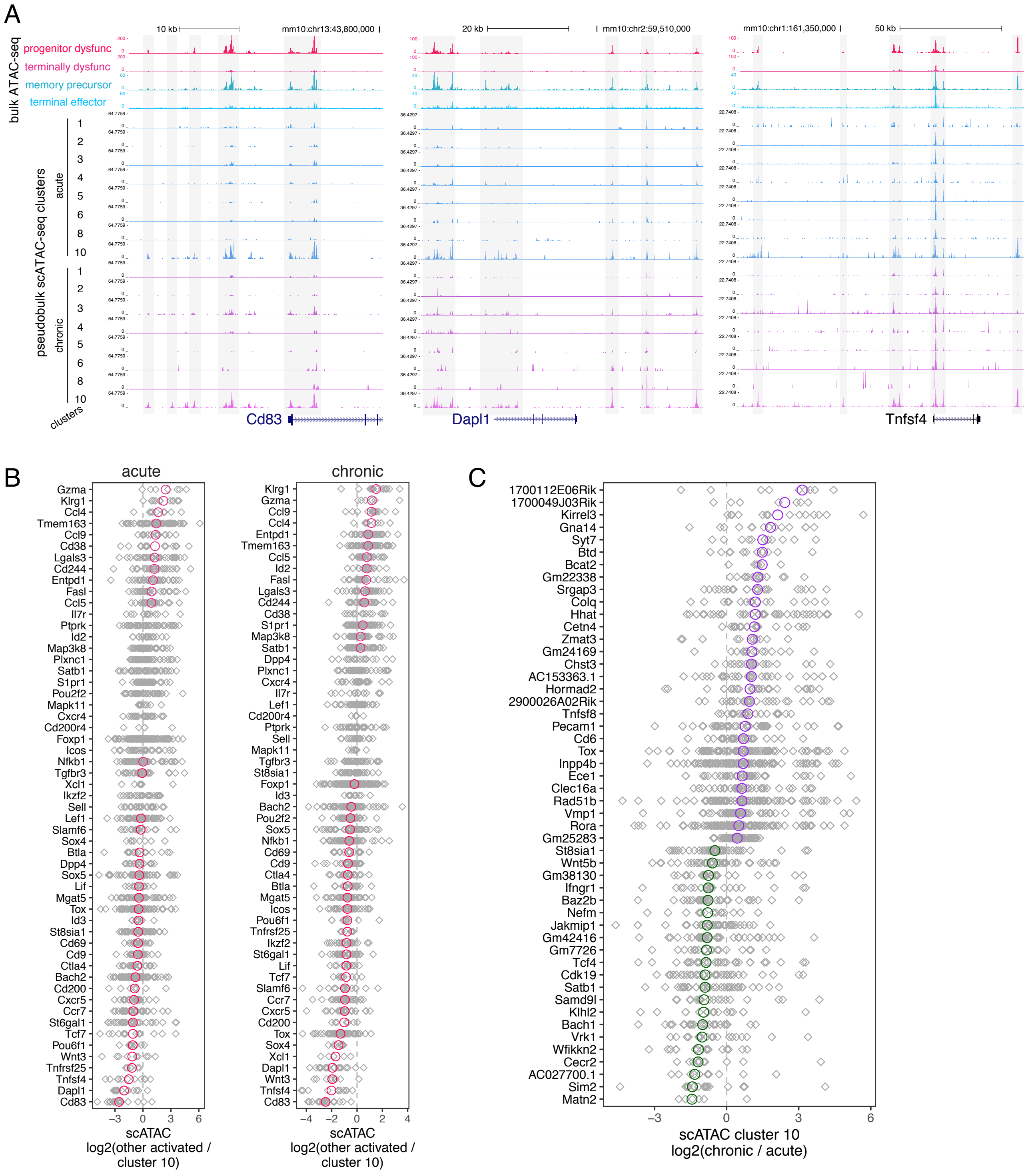

#### Supplementary Figure 8

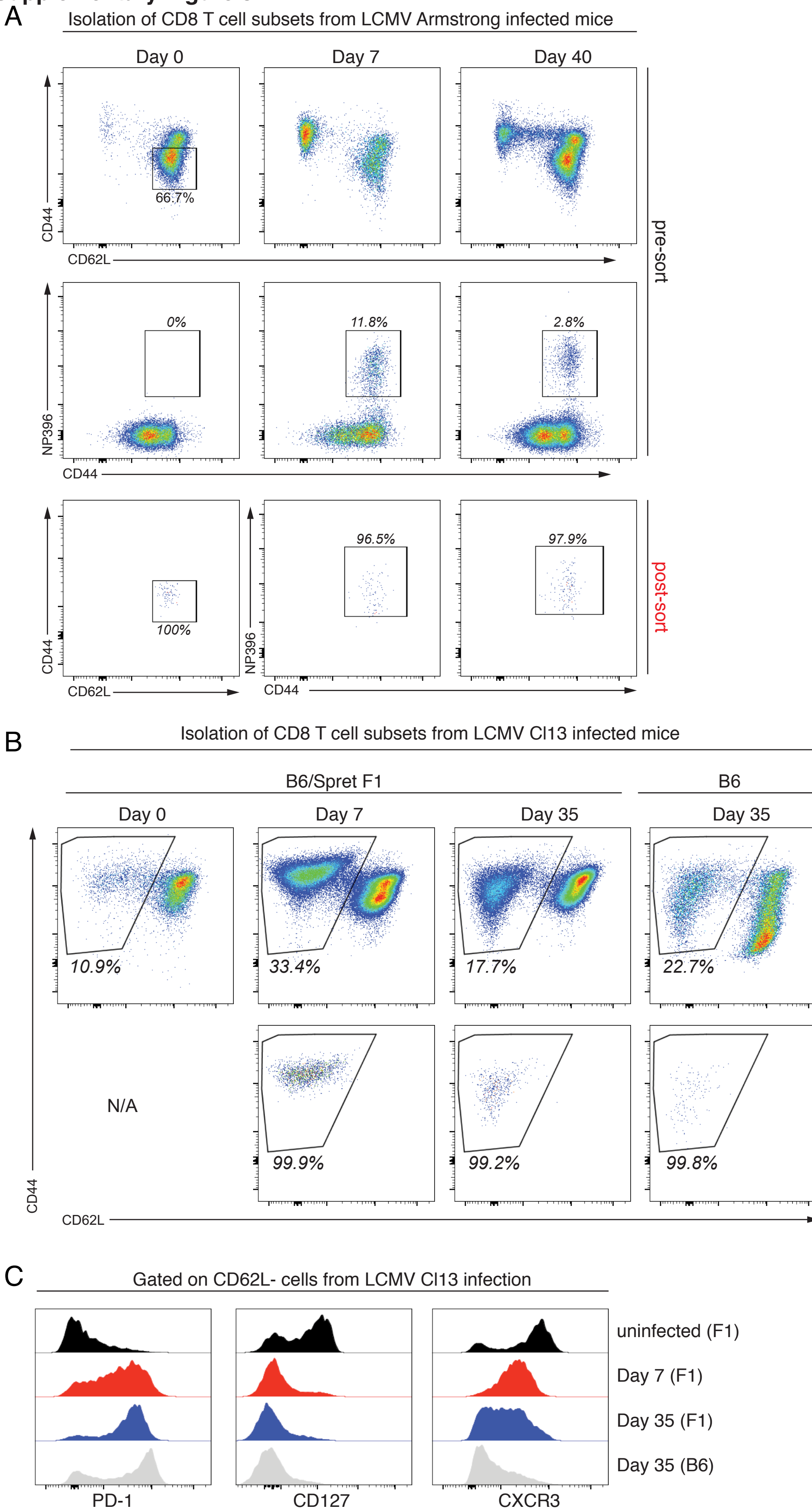

#### Supplementary Figure 9

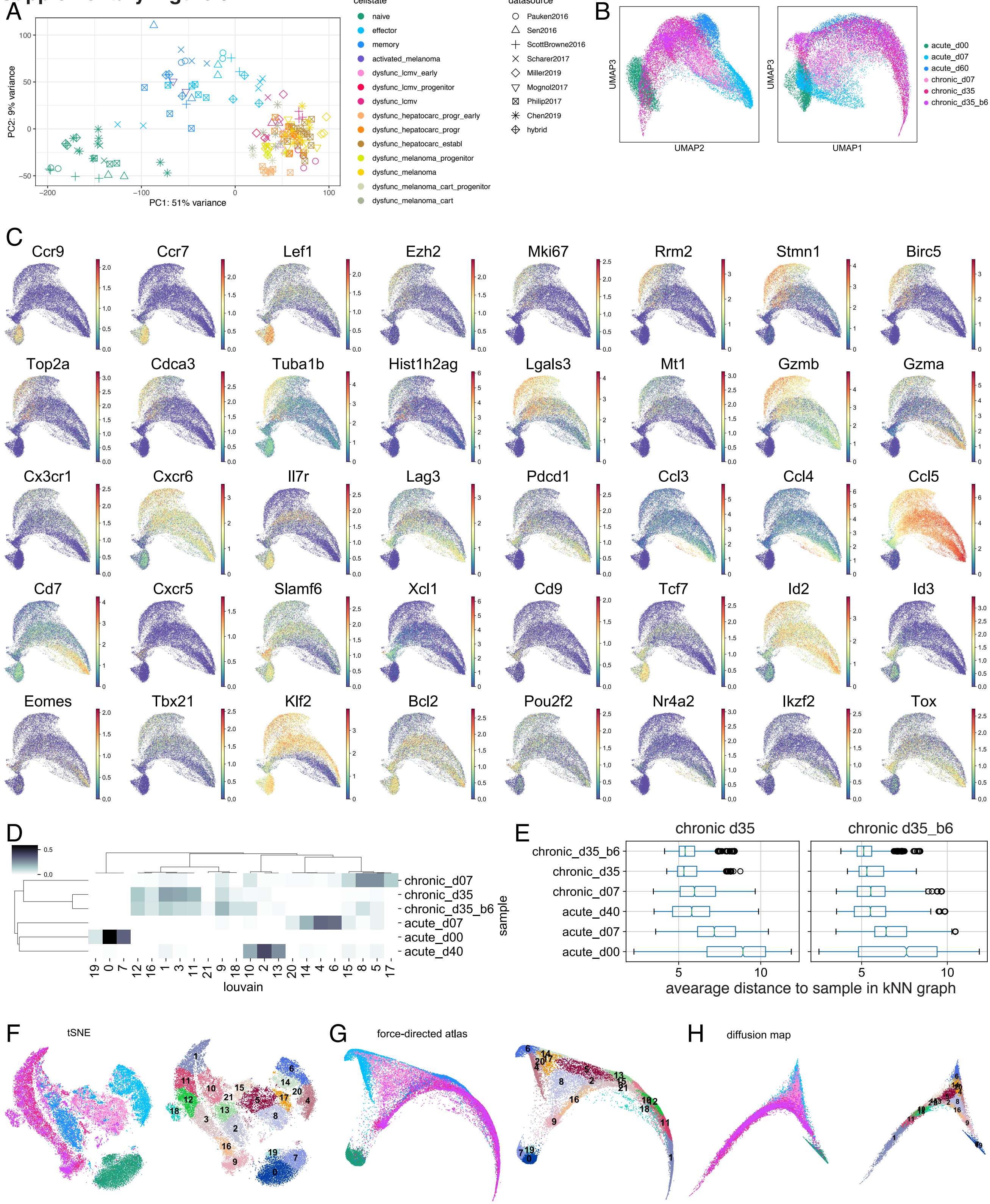

### Supplementary Figure 10

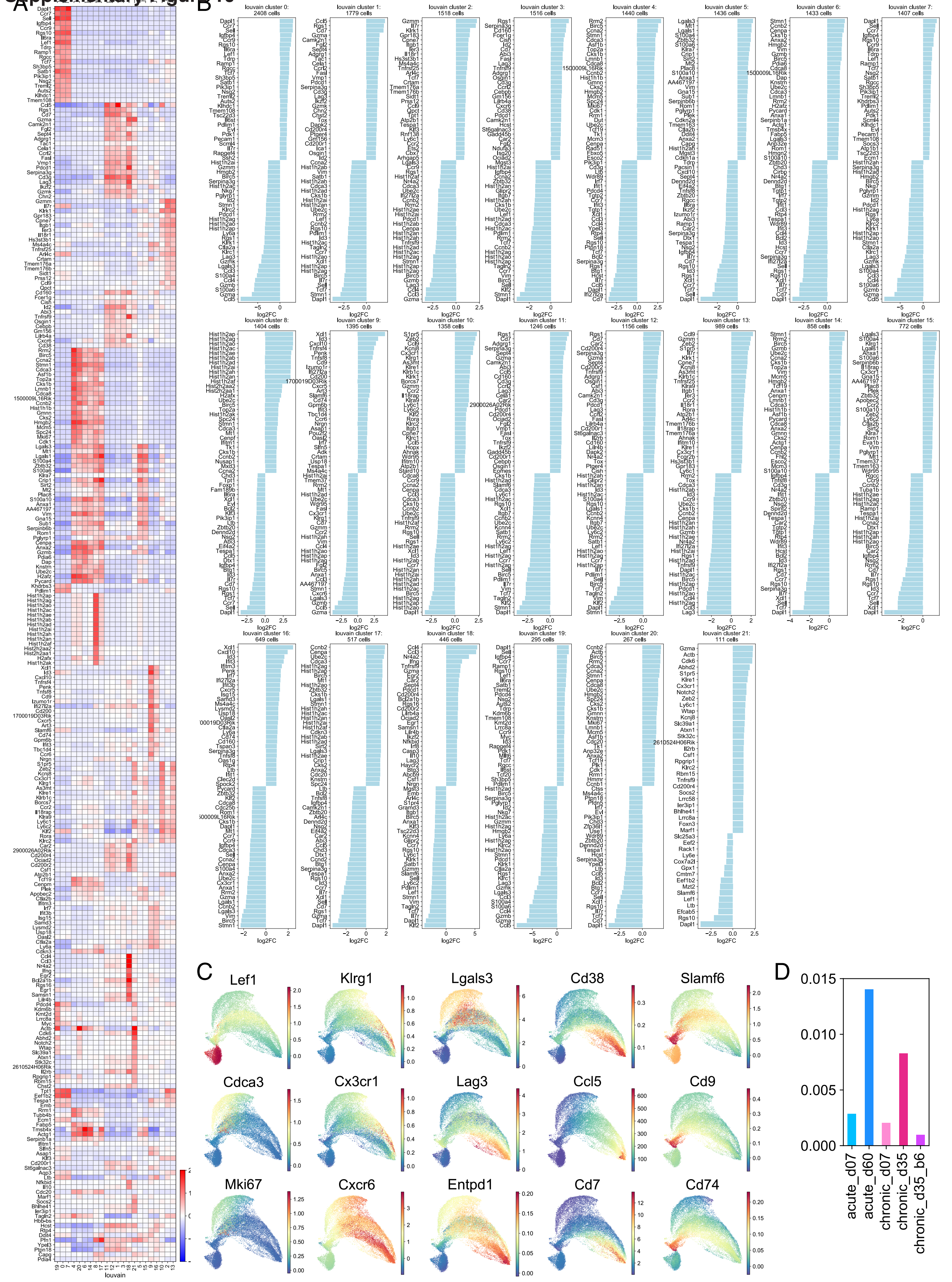

### Supplementary Figure 11

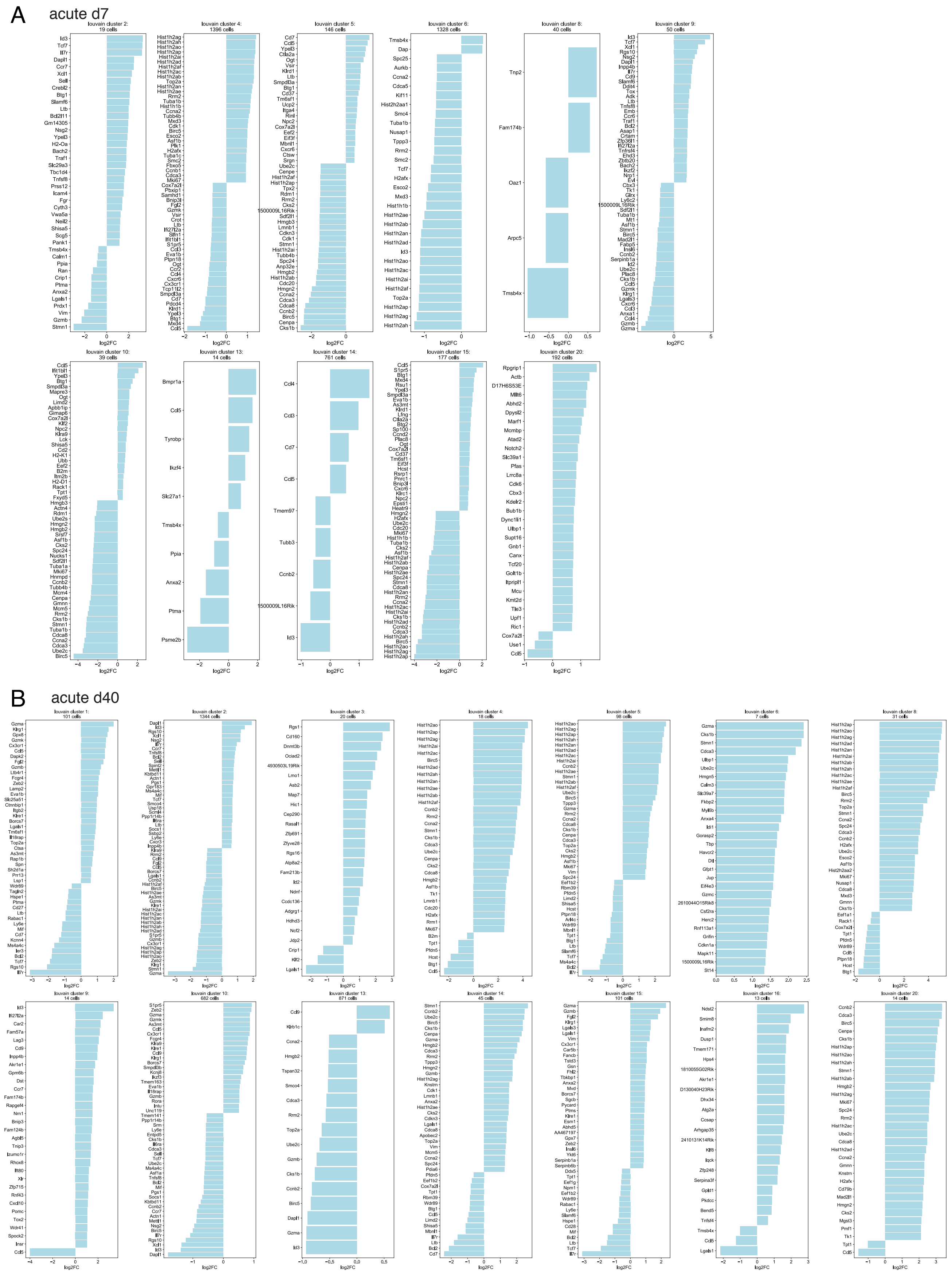

### Supplementary Figure 11

C chronic d7

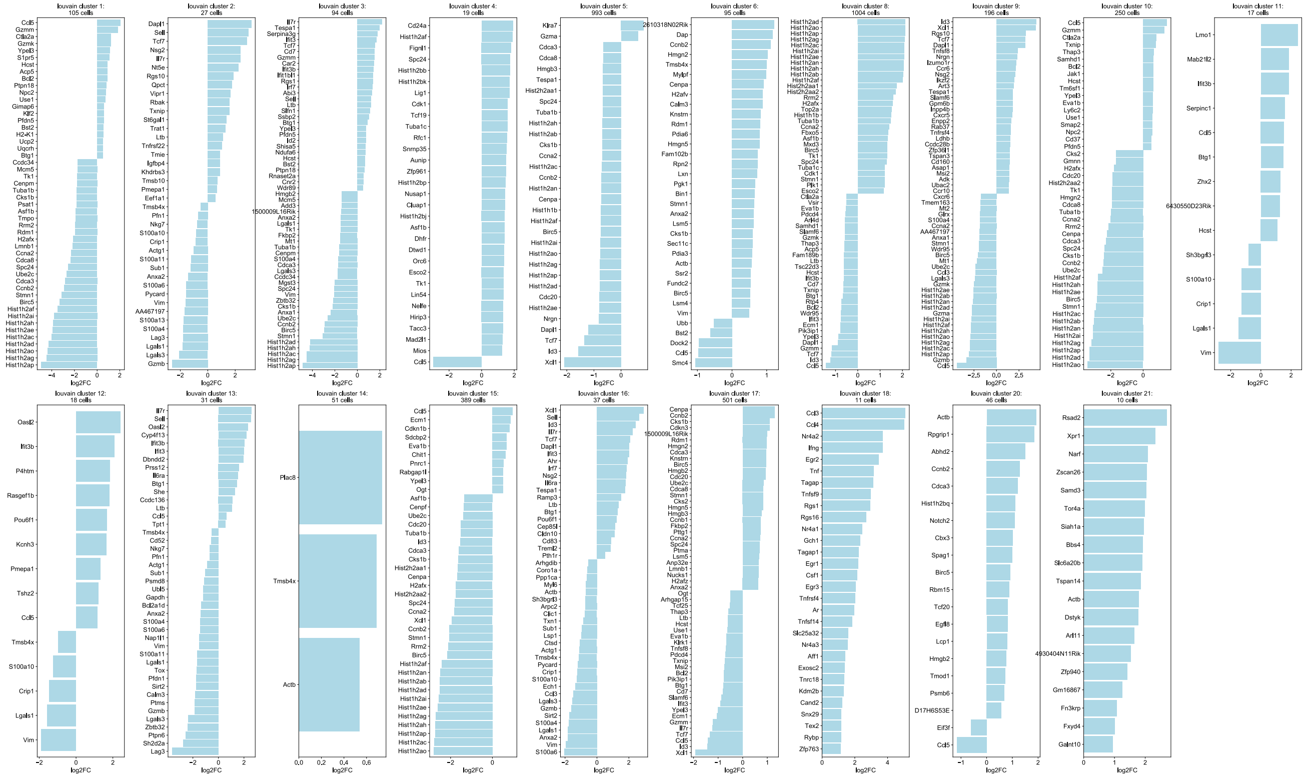

D chronic d35

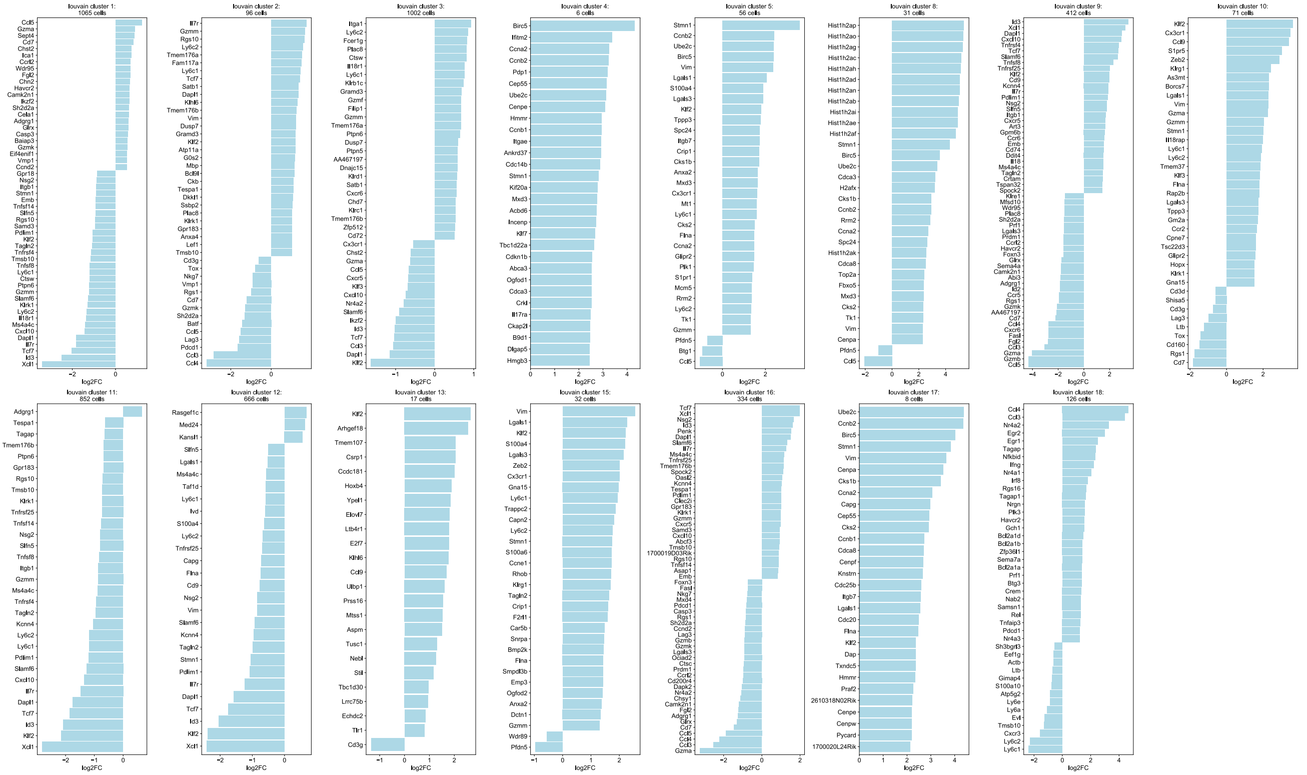

### Supplementary Figure 12

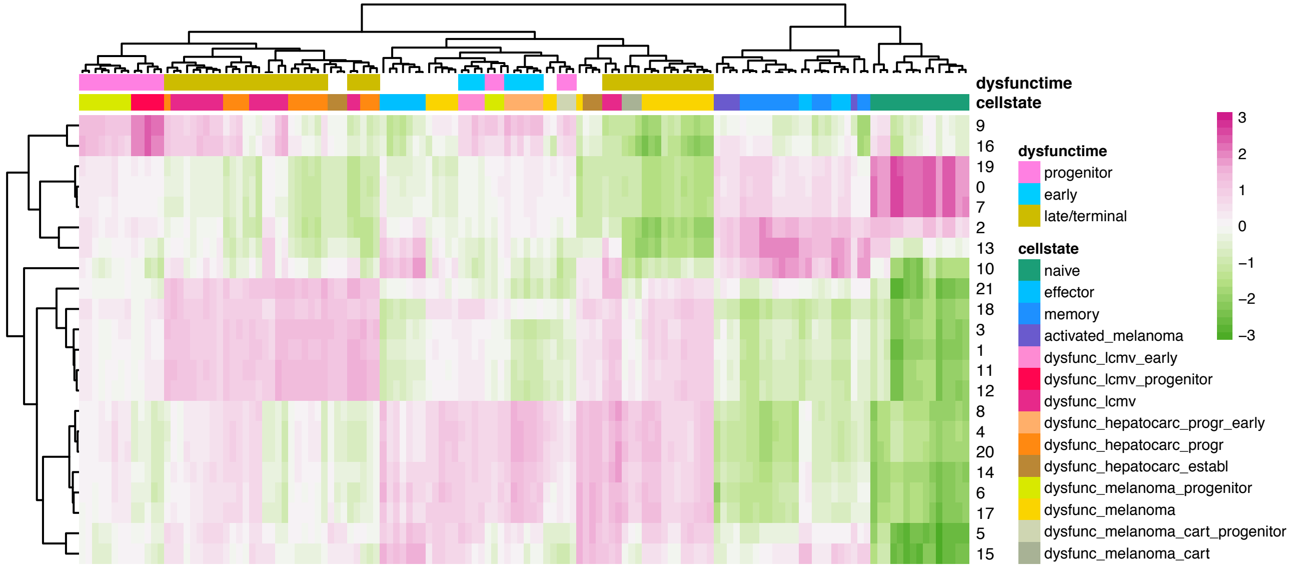

### Supplementary Figure 13

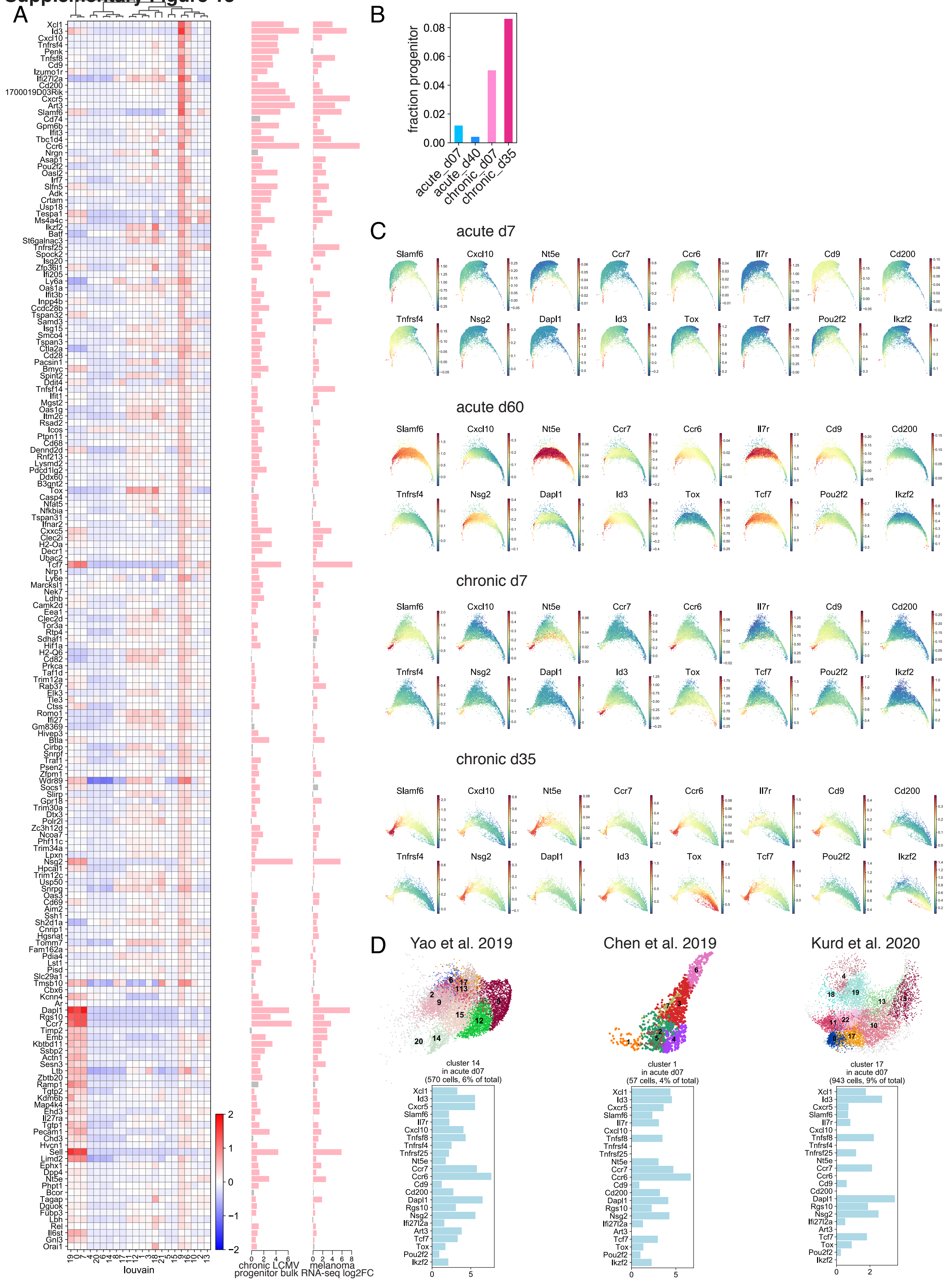

Supplementary Figure 14

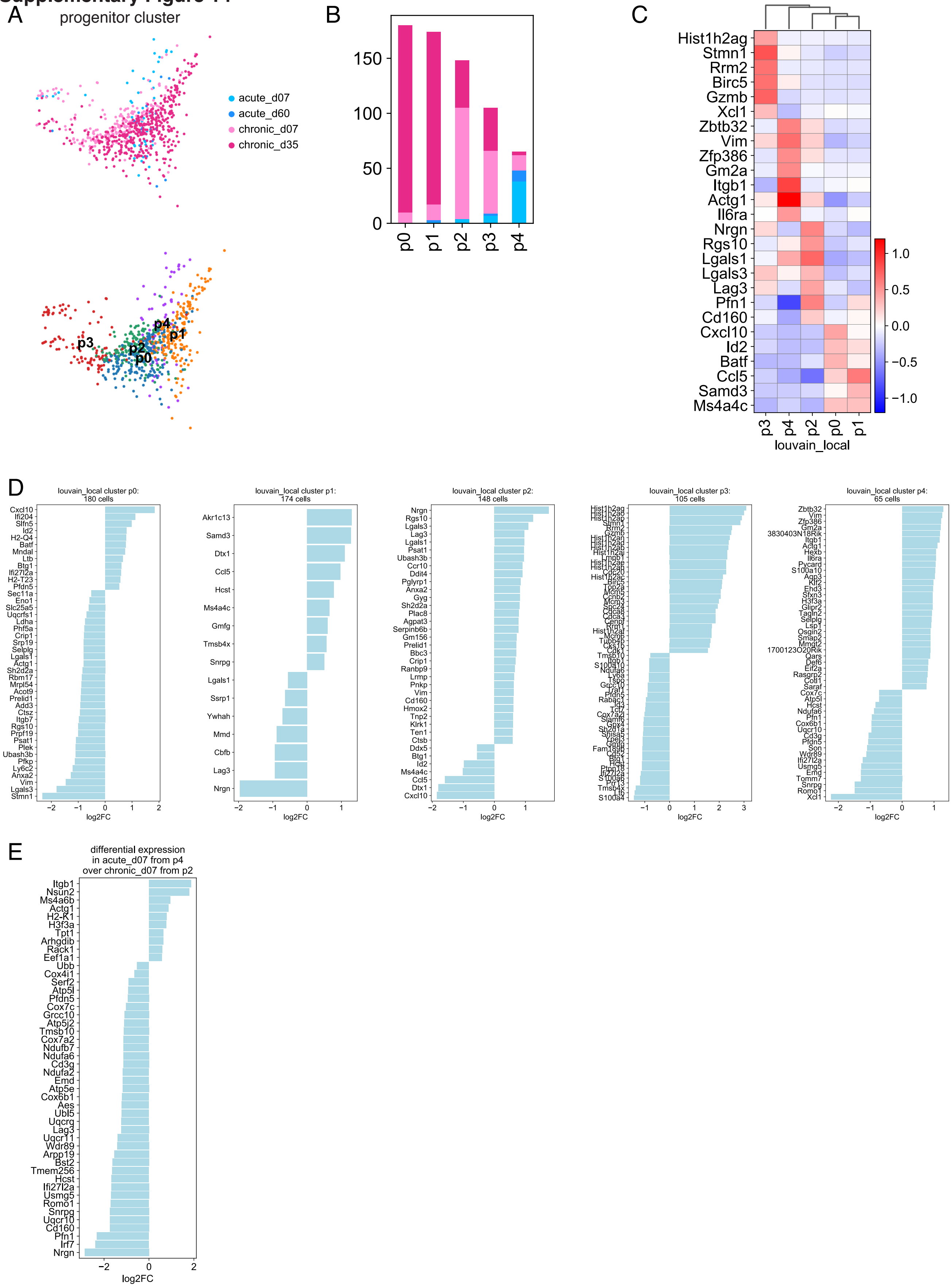

#### Supplementary Figure 15

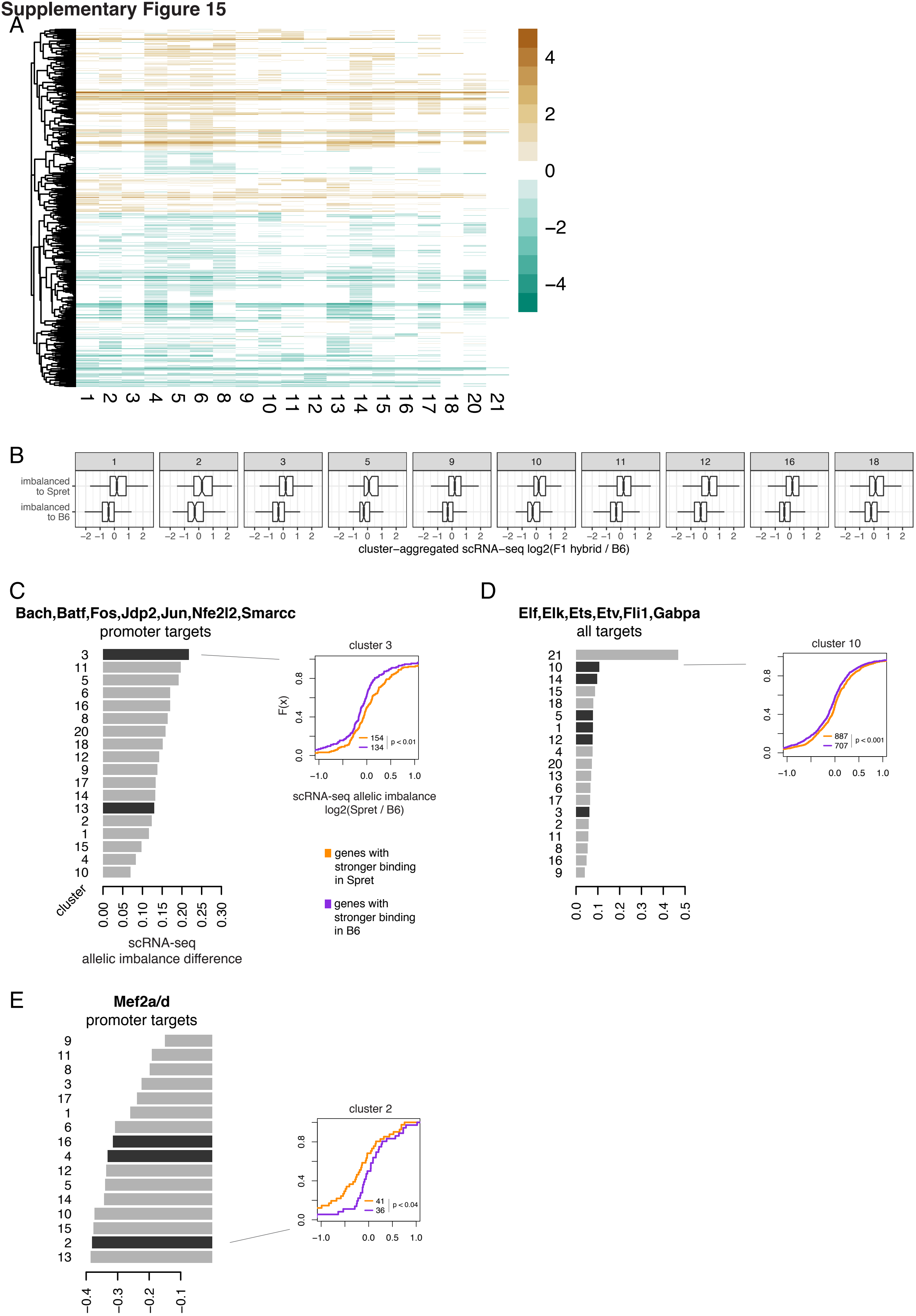

### Supplementary Figure 16

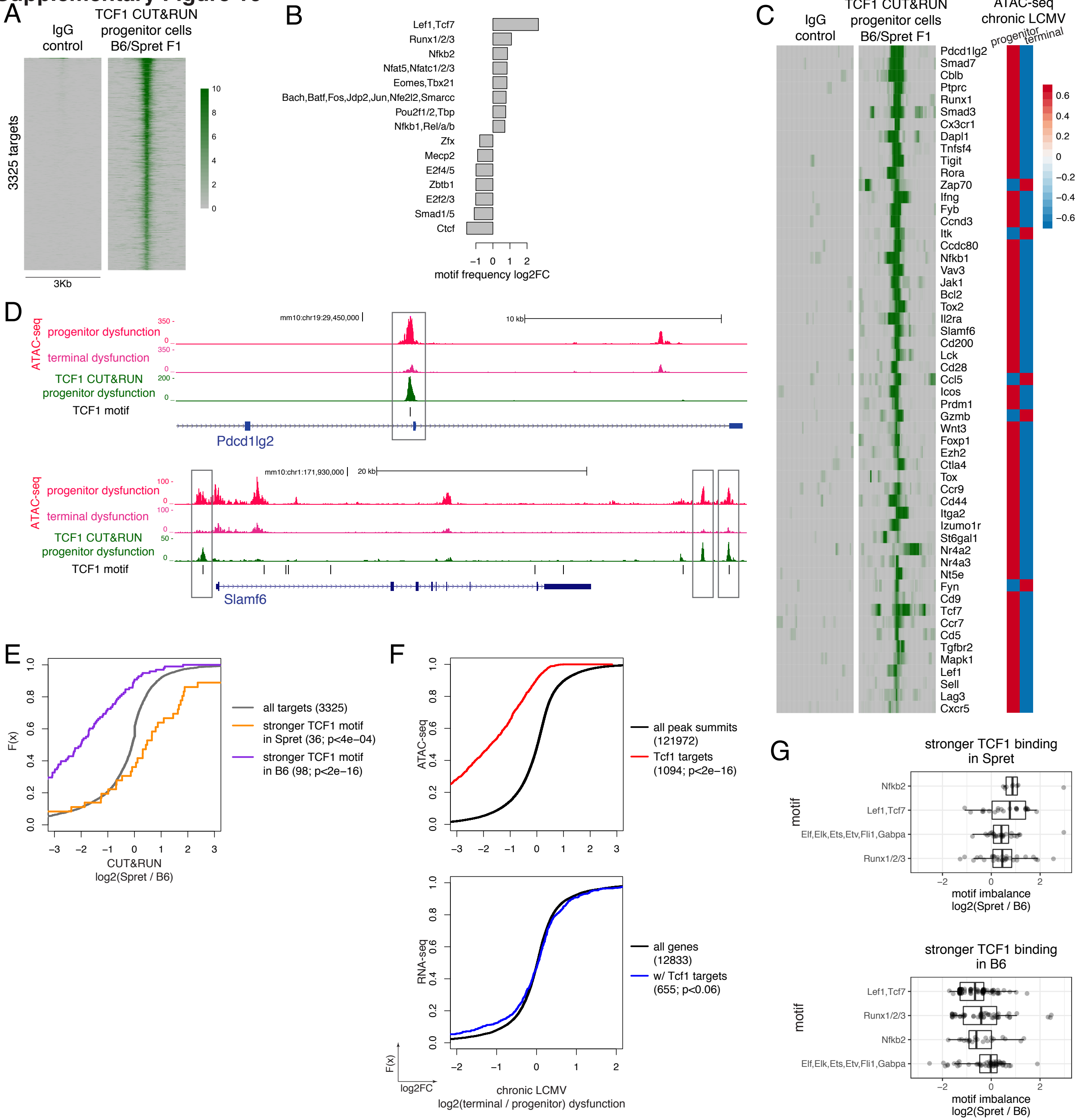

### Supplementary Figure 17

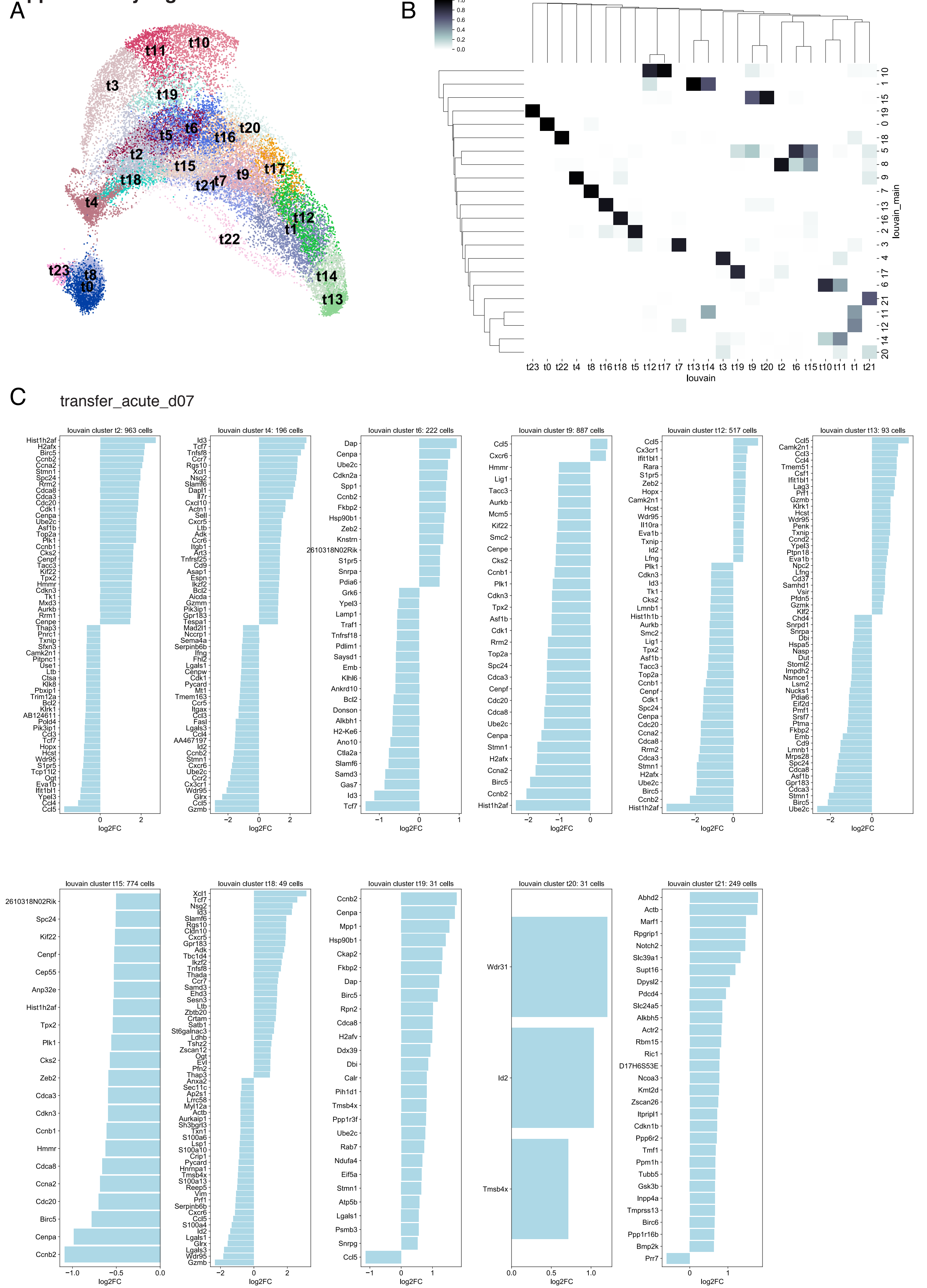
